# What executive function network is that? An image-based meta-analysis of network labels

**DOI:** 10.1101/2020.07.14.201202

**Authors:** Suzanne T. Witt, Helene van Ettinger-Veenstra, Taylor Salo, Michael C. Riedel, Angela R. Laird

**Affiliations:** BrainsCAN, University of Western Ontario, London, ON N6A 3K7, CANADA; Center for Social and Affective Neuroscience (CSAN), Linköping University/US, 581 85 Linköping, Sweden; Center for Medical Image Science and Visualization (CMIV), Linköping University/US, 581 85 Linköping, Sweden; Department of Psychology, Florida International University, Miami, FL 33199, USA; Department of Physics, Florida International University, Miami, FL 33199, USA

**Author notes:** **Corresponding Author:** Suzanne T. Witt, BrainsCAN, Western Interdisciplinary Research Building (WIRB), Rm 3190, University of Western Ontario, London, ON N6A 3K7, CANADA. Both authors contributed equally (Suzanne T. Witt, Helene van Ettinger-Veenstra). **Declarations**. **Conflicts of Interest/Competing Interests:** The authors report no conflicts of interest or competing interests. **Availability of data and material:** Unthresholded and thresholded statistical maps of the image-based meta-analysis results are available for download from NeuroVault (https://identifiers.org/neurovault.collection:8448). **Code availability:** The NiMARE software used for the image-based meta-analysis results reported in this manuscript is available for download (https://nimare.readthedocs.io/en/latest/index.html; https://github.com/neurostuff/NiMARE). **Author Contributions:** HvEV, STW, MCR, and ARL conceived of and designed the study. HvEV and STW performed the literature search and collected the published SPMs. TS and MCR contributed scripts, performed meta-analyses, and drafted the figures. HvEV and STW co-wrote the manuscript. All authors contributed to the revisions and approved the final version. HvEV and STW contributed equally to this work.

**Keywords:** executive function, brain networks, fMRI, image-based meta-analysis, network labels

## Abstract

The current state of label conventions used to describe brain networks related to executive functions is highly inconsistent, leading to confusion among researchers regarding network labels. Visually similar networks are referred to by different labels, yet these same labels are used to distinguish networks within studies. We performed a literature review of fMRI studies and identified nine frequently-used labels that are used to describe topographically or functionally similar neural networks: central executive network (CEN), cognitive control network (CCN), dorsal attention network (DAN), executive control network (ECN), executive network (EN), frontoparietal network (FPN), working memory network (WMN), task positive network (TPN), and ventral attention network (VAN). Our aim was to meta-analytically determine consistency of network topography within and across these labels. We hypothesized finding considerable overlap in the spatial topography among the neural networks associated with these labels. An image-based meta-analysis was performed on 158 group-level statistical maps (SPMs) received from authors of 69 papers listed on PubMed. Our results indicated that there was very little consistency in the SPMs labeled with a given network name. We identified four clusters of SPMs representing four spatially distinct executive function networks. We provide recommendations regarding label nomenclature and propose that authors looking to assign labels to executive function networks adopt this template set for labeling networks.

## Introduction

Functional neuroimaging studies frequently report connectivity or activation results within executive control networks that recruit frontoparietal areas, including the dorsolateral prefrontal and lateral posterior parietal regions. However, the current state of naming conventions for these commonly observed functional brain patterns is inconsistent throughout the literature, which limits a researcher’s ability to compare patterns across studies using network labels alone. An array of author-specified labels can be used to describe frontoparietal patterns, with no community consensus as to how these labels are defined. This includes, but is not limited to: the central executive network (CEN), cognitive control network (CCN), dorsal attention network (DAN), executive control network (ECN), executive network (EN), frontoparietal network (FPN), frontoparietal control network (FPCN), working memory network (WMN), task positive network (TPN), and ventral attention network (VAN). While these networks can appear topographically similar across studies, often they may differ in nuanced yet meaningful ways. It is especially confusing when these terms are differentially used within the same study to refer to topographically dissimilar networks. Related, but potentially separable, executive, control, or attentional mechanisms are likely associated with the published networks described by these labels. However, any attempt to derive a clear understanding of different topographical networks associated with different executive functions from the literature is not possible due to the widespread inconsistency in overlap or separation of names assigned to relevant topographic distributions.

There is community interest in developing a common nomenclature of functional brain networks. Many authors use published templates derived from large, canonical studies for automated or manual spatial sorting of networks and label assignment. Visual inspection of the published templates of the seminal studies from Smith et al. (Smith et al. 2009) and Yeo et al. (Yeo et al. 2011) reveals that the ECN (Fig. 1a), DAN (Fig. 1b), and left and right FPN (Fig. 1c) appear to be distinct, non-overlapping networks. However, even a cursory glance at the literature suggests that studies are using many of the different labels mentioned above to identify networks that are visually similar to these published templates.

**Figure 1.**
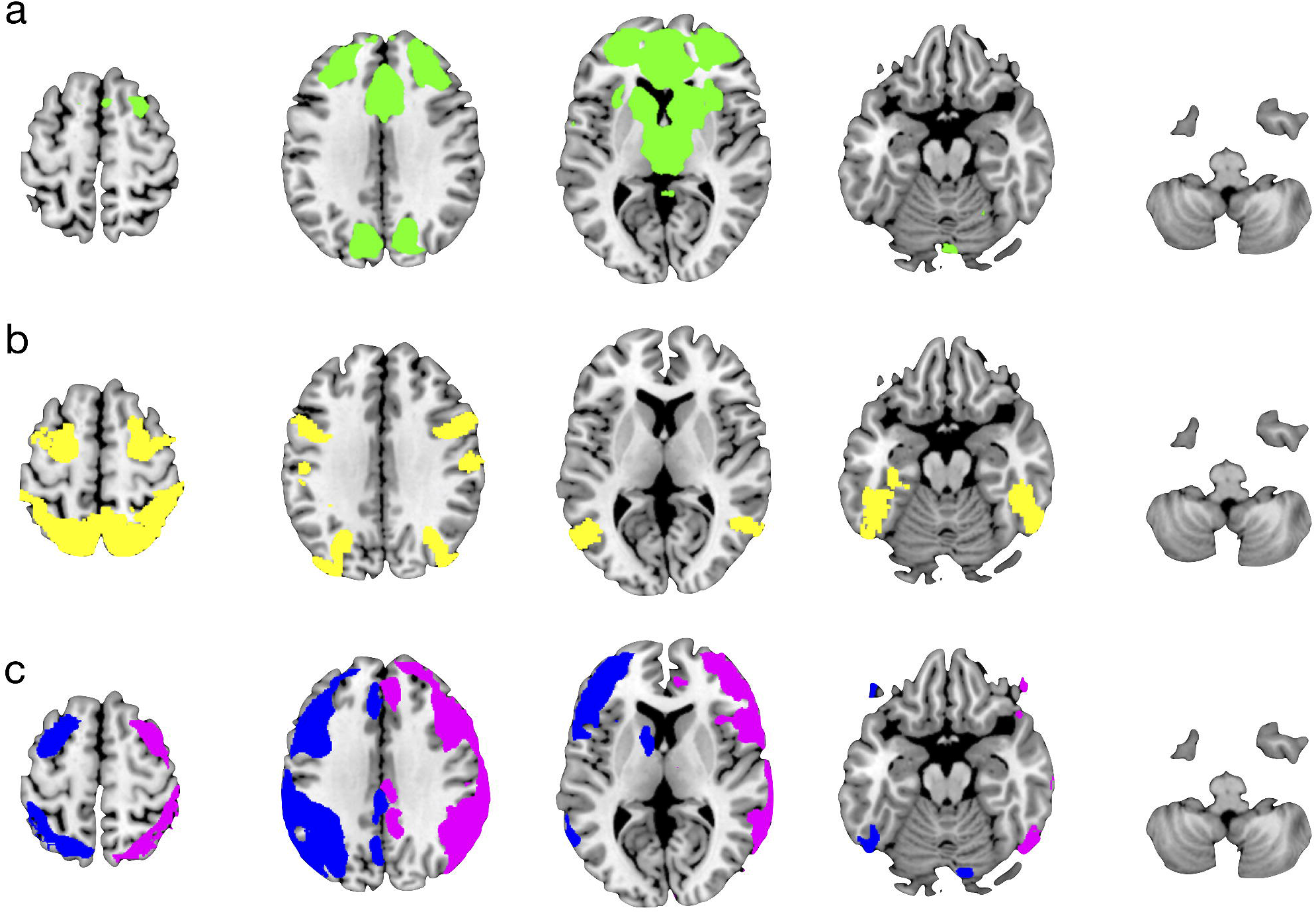
Existing templates of executive networks. a) Executive Control Network (ECN) from Smith et al. (Smith et al. 2009). b) Dorsal Attention Network (DAN) from Yeo et al. (Yeo et al. 2011). c) Left and Right Fronto-Parietal Networks (FPN) from Smith et al. (Smith et al. 2009).

The lack of consensus and inconsistencies in defining frontoparietal patterns in the literature may hinder the fMRI community’s efforts to discuss and interpret our collective findings. Here, we sought to explore whether the labels related to executive control consistently overlap with or are clearly separable from different topographical brain networks. To this end, we performed an image-based meta-analysis of group-level statistical parametric maps reported in the fMRI literature. The two extreme solutions to the label-network mapping consist of either one single topographical network to which all labels refer, or a one-to-one mapping between all possible topographical patterns and labels. Both extremes are unlikely; thus, we predict that the solution lies somewhere in the middle. Studies such as Smith et al., (Smith et al. 2009) and Yeo et al., (Yeo et al. 2011) have identified multiple executive control networks which are clearly topographically separable. Moreover, Laird et al. (Laird et al. 2011) leveraged metadata from the BrainMap database to demonstrate that multiple frontoparietal networks can be mapped to separate mental functions. Additionally, topographically similar networks identified by multiple, different network labels suggests that networks and labels do not map one-to-one. We hypothesize that multiple separable patterns with dissociable spatial topographies exist, are commonly reported in the literature, and yet are not reflected in our current collective strategy for naming executive control networks. To test our hypothesis, two separate analyses were performed. The first analysis aimed to assess the within- and across-network consistency of commonly used network labels to confirm that a one-to-one mapping between network label and topographically distinct networks does not exist. The second analysis aimed to identify underlying topographically separable networks in previously published group-level statistical parametric maps (SPMs) to determine the true number of neural networks related to executive functioning. Confirmation of this hypothesis would serve as a clear illustration of the lack of community consensus and the need for community-endorsed labeling recommendations.

## Methods

### Literature Search

PubMed was searched for fMRI studies that reported group-level SPMs explicitly labeled with any one of the network names of interest. Our search terms included, “fMRI central executive network”, “fMRI ‘central executive network’”, “fMRI cognitive control network”, “fMRI ‘cognitive control network’”, “fMRI dorsal attention network”, “fMRI ‘dorsal attention network’”, “fMRI executive network”, “fMRI ‘executive network’”, “fMRI executive control network”, “fMRI ‘executive control network’”, “fMRI frontoparietal control network”, “fMRI ‘fronto-parietal control network’”, “fMRI frontoparietal network”, “fMRI ‘frontoparietal network’”, “fMRI task positive network”, “fMRI ‘task positive network’”, “fMRI ventral attention network”, “fMRI ‘ventral attention network’”, “fMRI working memory network”, “fMRI ‘working memory network’”. Additional searches were performed for “fMRI ICA” and “fMRI ‘independent component analysis’” to capture studies not otherwise related to a specific network name. Search results were limited to include papers published over a period of 10 years (2007-2017), reporting results from adult participants aged nineteen years and older. These searches netted 7,864 results, including duplicates.

Papers were then individually examined to determine whether they met the following inclusion criteria: 1) the study included healthy participants of mean age no younger than 18 years and no older than 70 years; 2) the study presented statistical maps of healthy participants only or separately from patients; 3) statistical maps were derived from whole-brain analytic methods; and 4) statistical maps were explicitly defined with one of the targeted network names in the Methods, Results, Figure/Figure Captions, or Discussion section of the paper. Papers were excluded if they did not meet all of the inclusion criteria. Papers were also excluded if networks were defined using either seed-voxel or ROI based techniques.

Corresponding authors for all identified papers were emailed to request the group-level SPM file corresponding to the relevant network-labeled figure. If the email address for the corresponding author was no longer valid, the email request would be sent to the first or last author on the paper, along with a request for the best person to contact regarding the statistical maps. In cases where multiple maps were being requested from the same paper, a single email request was sent for all statistical parametric map files. Approximately three months after the initial email request, all studies that either responded requesting more time to search for the relevant files or had not responded to the initial email request were sent a second email request.

### SPM Preprocessing

The group-level SPMs netted from the search were each aligned to MNI space using FSL’s FLIRT toolbox (Jenkinson et al. 2002). Registration quality was assessed by overlapping the SPM on the 2-mm resolution MNI template, and observing agreement between between the brain boundaries and mask or ventricle edges in the SPMs. In a few instances, manual 3-parameter translations were performed using Mango to correct for alignment issues. Maps already normalized to MNI space were resliced to 2mm isotropic resolution using Mango (http://ric.uthscsa.edu/mango). Study-specific pre-processing pipelines results in different total brain sizes between SPMs; thus, a conservative masking approach was used in which only those voxels that contained a non-zero value across all SPMs were considered for further analysis. Each group-level SPM was then imported into MATLAB (Natick, MA) using SPM12. All meta-analyses were performed on unthresholded SPMs.

### Image-Based Meta-Analysis 1: Consistency Within and Across Labels

For the first analysis examining the consistency of the group-level SPMs within and across network labels, the SPMs were first separately variance normalized. To assess within-network label consistency of the spatial topographies, the SPMs assigned to each network label were correlated with one another. To assess the across-network label consistency of the spatial topographies, all SPMs were first averaged to generate an overall SPM of executive control. Then, all group-level SPMs assigned to each network label were averaged, and Pearson’s correlation coefficients were calculated between each average network label map and the map of overall executive control. In line with the prevailing interpretations of correlation coefficients, we considered a correlation coefficient greater than 0.6 to be indicative of a strong spatial correlation. Multiple comparisons were corrected using the Benjamini-Hochberg false-discovery rate (Benjamini and Hochberg 1995). To assess the similarity between network labels themselves, the average SPM images for each network label were subjected to a hierarchical clustering analysis using the Euclidean distance metric and Ward’s linkage algorithm.

### Image-Based Meta-Analysis 2: Spatial Topography of Distinct Networks

For the second analysis examining the underlying spatial topography of distinct networks contained within the group-level SPMs, the SPMs were transformed into 1D arrays and concatenated. The resulting voxel by input image matrix was subjected to a hierarchical clustering analysis using the Euclidean distance metric and Ward’s linkage algorithm. Based on visual inspection of the dendrogram, four clusters of SPMs were identified. To produce statistical maps for these four clusters, permutation-based, image-based meta-analyses were then performed for each of the four clusters using the NiMARE software package [RRID:SCR_017398] (Salo et al. 2018), where the SPMs assigned to each cluster were first variance-normalized and subjected to a one-sample t-test. A permutation-based null distribution was constructed for each voxel by randomly setting voxel-value signs based on the proportion of positive and negative values across the sample and performing a one-sample t-test on the randomized data. The distribution of t-values for each voxel was then used as the null distribution to assess the significance of each voxel’s original t-value. Multiple comparisons were corrected using the Benjamini-Hochberg false-discovery rate (Benjamini and Hochberg 1995).

## Results

### Literature Search

The literature search procedures resulted in the identification of 188 papers that met our defined search criteria; these studies reported results for 365 group-level statistical maps. Of the 365 statistical maps: 26 reported networks labeled as CEN, 13 CCN, 64 DAN, 46 ECN, 19 EN, 8 FPCN, 127 FPN, 19 TPN, 21 VAN, and 22 WMN. As detailed in Fig. 2, of the 188 papers identified during the initial PubMed search, 72 authors responded positively and sent the 166 requested SPMs, 21 responded negatively, 12 responded initially but did not respond to follow up emails, and the authors for the remaining 83 studies did not respond at all. Regarding the 21 papers that responded negatively, the most common reason for the negative response was lack of access to the data due to data being lost as a result of technical upgrades and/or failures. The literature search procedures yielded a total set of 166 group-level SPMs received from authors, which included 5 CEN, 7 CCN, 26 DAN, 21 ECN, 10 EN, 73 FPN, 2 FPCN, 7 TPN, 7 VAN, and 8 WMN maps. As only two SPMs labeled FPCN were received, we chose to group the SPMs labeled FPN and FPCN into a single group, under the label FPN.

**Figure 2.**
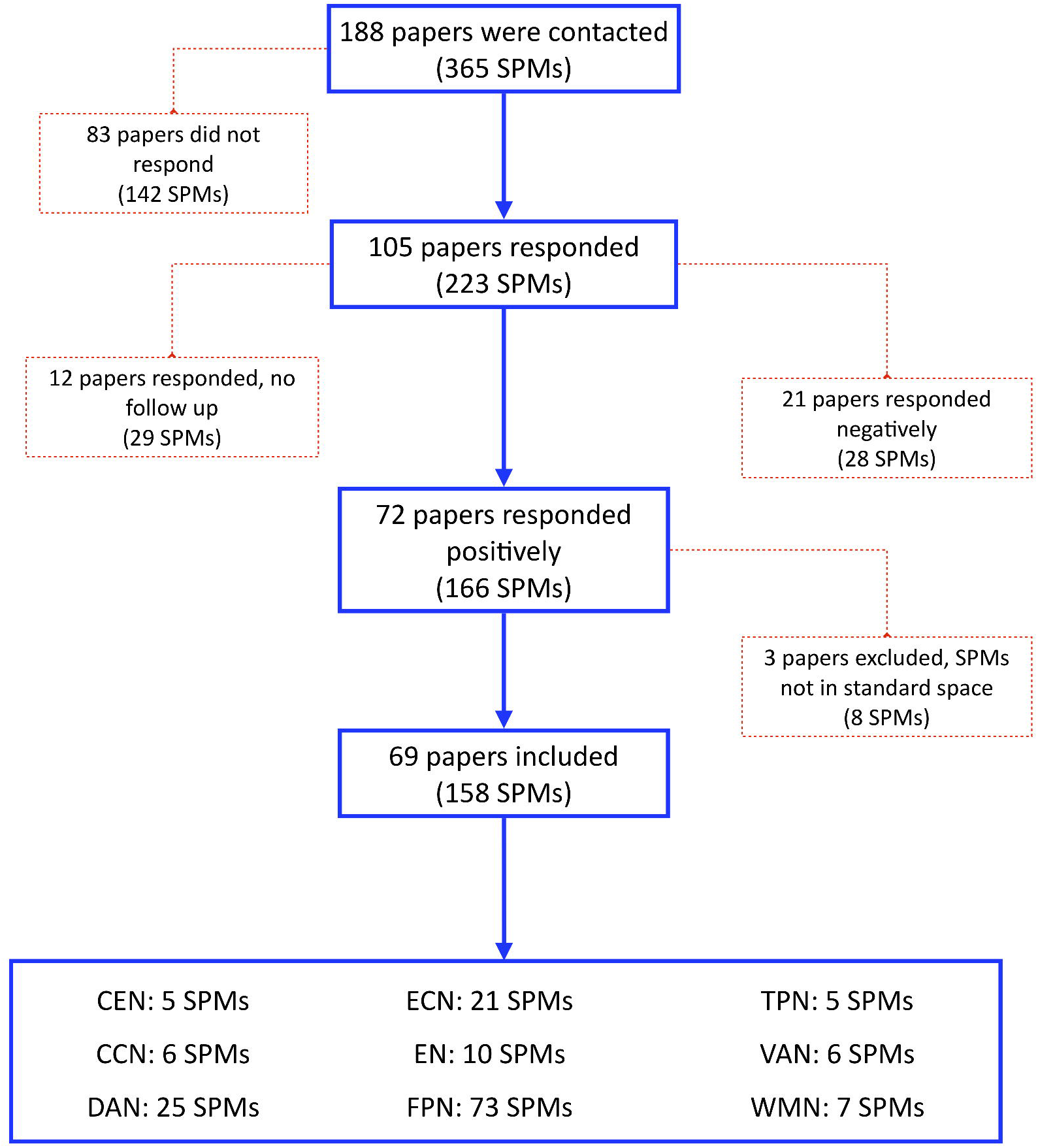
Flowchart of literature search results. The blue path with solid lines and arrows indicates positive search results where authors sent the requested statistical parametric maps (SPMs) and the received SPMs passed all quality checks. The red, dashed path indicates negative search results reflecting when authors did not respond to either initial or follow-up requests, failed to send requested SPMs, were not able to send requested SPMs, or the SPMs failed quality checks. Network labels refer to the central executive network (CEN), cognitive control network (CCN), dorsal attention network (DAN), executive control network (ECN), executive network (EN), frontoparietal network (FPN), frontoparietal control network (FPCN), working memory network (WMN), task positive network (TPN), and ventral attention network (VAN).

### Final Sample of Group-Level SPMs

The 166 group-level SPMs received from authors were individually examined to ensure the image files could be successfully loaded and met the study inclusion criteria. Of those, eight SPMs from three studies were discarded due to the maps not being in either MNI or Talairach space. No SPMs were discarded due to file corruption. These quality checks resulted in a final sample of 158 group-level SPMs (5 CEN, 6 CCN, 25 DAN, 21 ECN, 10 EN, 73 FPN, 5 TPN, 6 VAN, 7 WMN) from 69 studies (Fig. 2). Of these 69 studies, 37 reported results for resting-state data and 31 for task-based data. One study reported results for both resting-state and task-based data to separately examine the same executive function networks. Task-based studies generally reported fewer SPMs per paper compared to resting-state studies, due to the prevalence of ICA-based studies reporting multiple component maps per paper. Thus, 51 group-level SPMs in our final sample were estimated from task-based data and the remaining 107 group-level SPMs from resting-state data. The majority of the task-based studies employed the use of traditional executive function tasks (n=26). These included working memory tasks, such as the n-back and the delayed match-to-sample tasks; cognitive control tasks, such as the Stroop, Flanker, and saccades tasks; and attention tasks, such as the revised attention network, continuous attention, and visual-spatial attention tasks. The remaining five task-based studies employed tasks engaging processes related to language, reward anticipation, autobiographical memory, and social communication. A more complete description of the final meta-analytic sample, including full citations and relevant study details, is included in Supplemental Table 1.

### Image-Based Meta-Analysis 1: Consistency Within and Across Labels

We examined the consistency of SPMs within network labels and observed that the correlation between any two group-level SPMs labeled with the same network label was less than 0.6 (Fig. 3a solid line). This suggests that there is an overall lack of consistency in the spatial topography of the neural networks described by any given network label. Further, we examined the consistency of spatial topography between network labels and the average executive control map of all SPMs (Fig. 3a dashed line) and observed that the average maps for EN, FPN, CEN, and WMN exhibited a correlation of greater than 0.8 with this overall SPM of executive control. The most extreme example of this pattern was seen with the CEN, for which there was virtually no spatial correlation within the five group-level SPMs using this label, but the average CEN map exhibited strong spatial correlation with the average executive control map. These dual findings for the CEN suggest that this label is being applied to individual maps of executive function with differing spatial topographies, but that the label itself could be used to identify any number of executive function related spatial topographies. This presence of both weak within-network correlation and strong between-network correlation confirmed our study hypothesis that multiple executive control network labels (i.e., EN, FPN, CEN, and WMN) are being used to describe topographically similar networks. The average maps for the labels VAN, ECN, TPN, CCN, and DAN, however, had a correlation of less than 0.6 with the overall SPM map of executive control, indicating that there are spatially distinct topographies of executive control networks.

**Figure 3.**
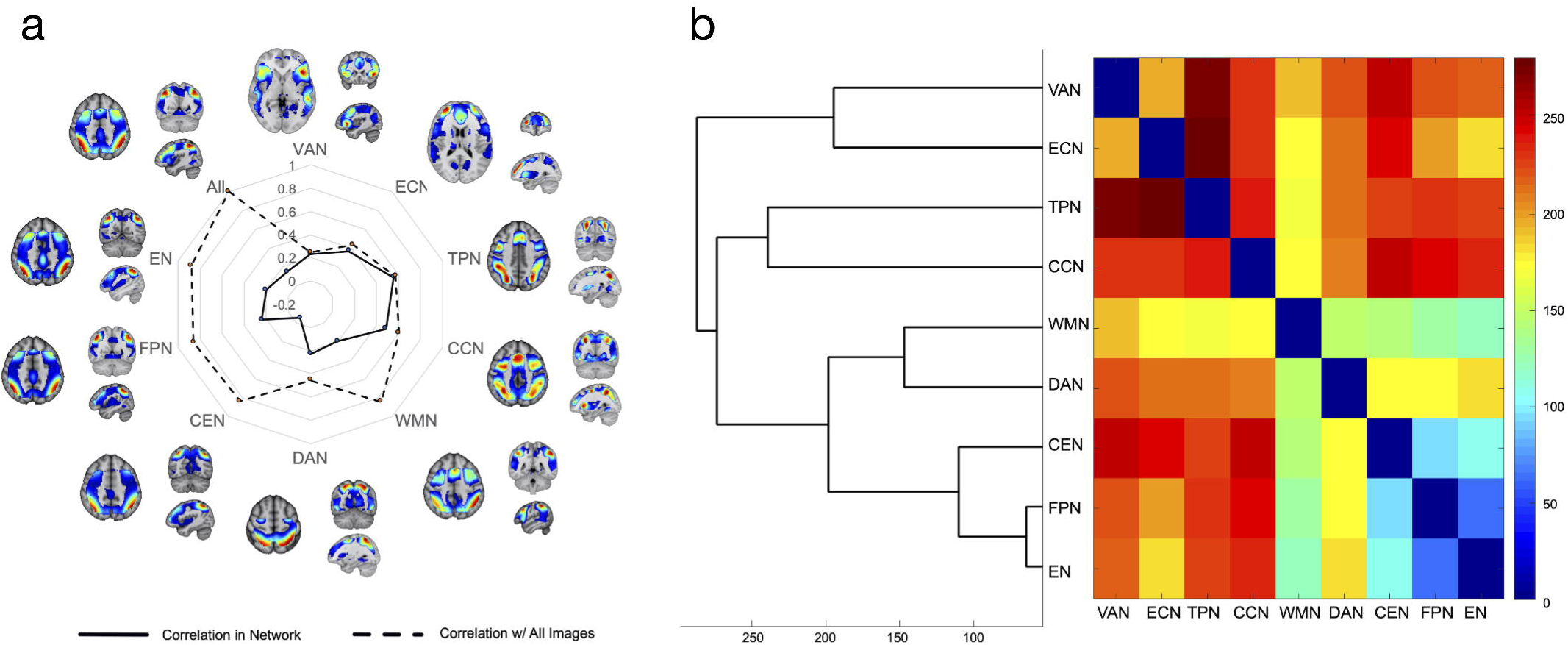
Consistency within and across labels. a) Results of spatial consistency among SPMs. The solid line indicates the average Pearson’s correlation coefficient between any two SPMs labeled with the same network label (i.e., within-network label consistency). The dashed line shows the Pearson’s correlation between the average of all SPMs for a given network label and all SPMs included in the analysis (across-network label consistency). b) Dendrogram showing the similarity among the nine network labels of interest. The horizontal axis of the dendrogram represents Ward’s linkage.

Hierarchical clustering of the average SPMs for the network labels yielded two major clusters (Fig. 3b). The first cluster consisted of the VAN and ECN labels, while the second cluster included the remaining seven network labels. This second main cluster was further subdivided into three label sub-clusters: 1) TPN and CCN, 2) WMN and DAN, and 3) CEN, FPN, and EN. The similarity of the network labels, as measured by Euclidian distance, paralleled the results described above for correlating the average SPM for a given network label with the overall SPM of executive control. Namely, the WMN, CEN, FPN, and EN networks exhibited the greatest similarity.

### Image-Based Meta-Analysis 2: Spatial Topography of Distinct Networks

The second clustering analysis to identify distinct spatial networks contained in the group-level SPMs resulted in four clusters of SPMs (Fig. 4a). Permutation-based, image-based meta-analysis was then performed for these four clusters to produce a topographic map for each. A more detailed visual depiction of the topographic maps produced by the permutation-based, image-based meta-analysis is shown in Supplemental Figure 1.

**Figure 4.**
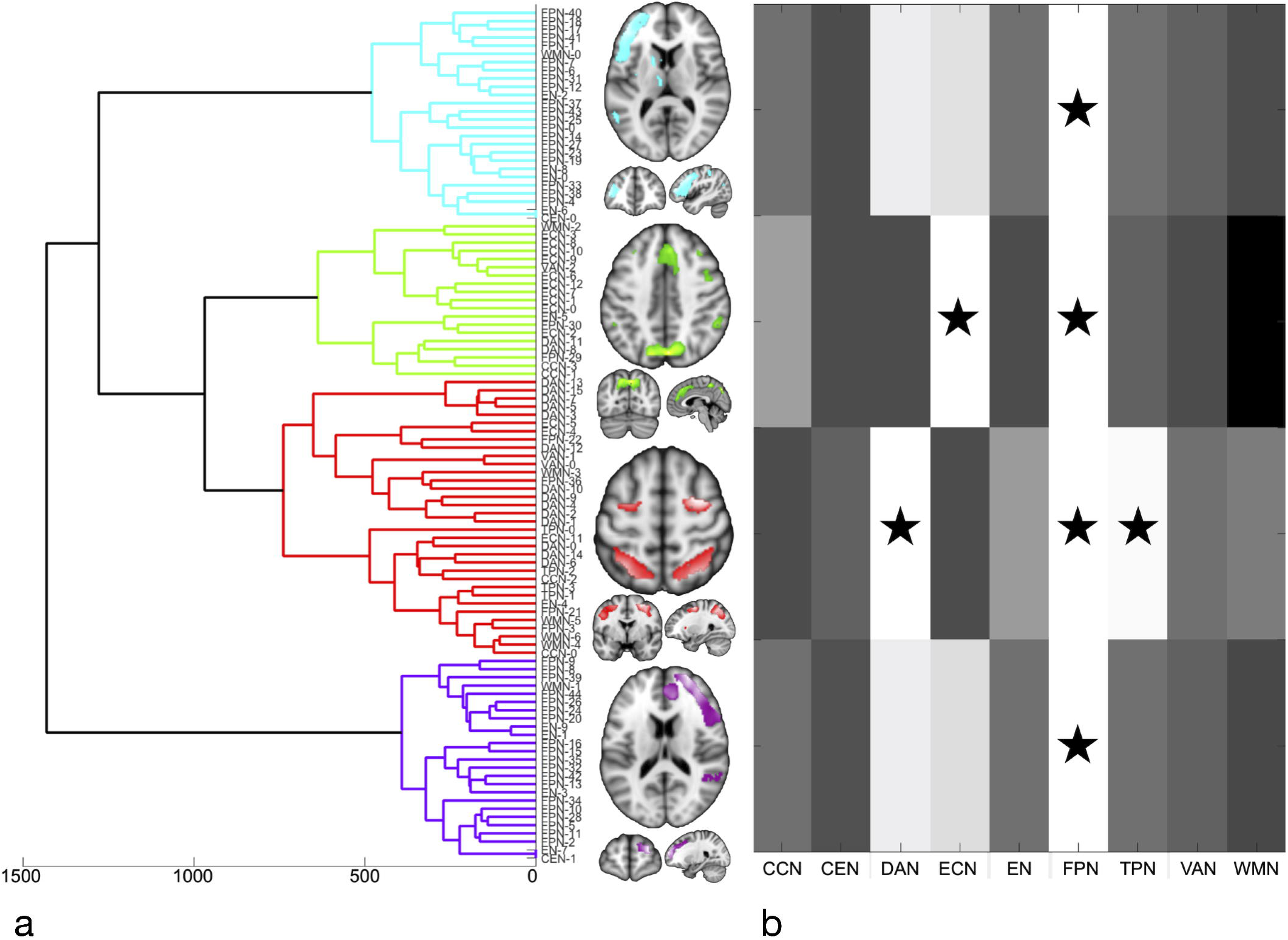
Spatial topography of distinct networks. a) Four separable network clusters were identified from the SPMs included in this analysis. b) SPMs labeled “fronto-parietal network” (FPN) were found to contribute to all four clusters. SPMs labeled “executive control network” (ECN) were found to contribute to the green cluster. SPMs labeled “dorsal attention network” (DAN) and “task positive network” (TPN) were found to contribute to the red cluster. None of the other five network labels investigated were found to significantly contribute to any of the four clusters.

### Cyan Cluster

The peak regions in the cyan cluster were primarily localized to the left hemisphere (Supplemental Fig. 2, Supplemental Table 2). The largest peak region was located in the left middle frontal gyrus. The next largest peak region encompassed the left precuneus and left superior and inferior parietal lobes. Additional peak regions in the cyan cluster were localized in left middle and inferior temporal gyri; left posterior cingulate gyrus; left caudate and claustrum; left thalamus; and left cingulate gyrus. Finally, two peak regions were found in the right hemisphere in right superior parietal lobe and right middle frontal gyrus.

### Green Cluster

The peak regions in the green cluster (Supplemental Fig. 3, Supplemental Table 3) were primarily located in bilateral medial frontal gyri; bilateral precuneus extending into paracentral lobule and superior parietal lobe; and bilateral middle and superior frontal gyri. Additional peak regions were found in left putamen and insula; bilateral precentral gyri; and bilateral inferior parietal lobes.

### Red Cluster

The peak regions in the red cluster (Supplemental Fig. 4, Supplemental Table 4) were primarily located in bilateral superior and inferior parietal lobes and bilateral frontal sub-gyral. Additional peak regions were observed in bilateral insulae and bilateral superior frontal gyri.

### Purple Cluster

The purple cluster (Supplemental Fig. 5, Supplemental Table 5) appeared to be a right hemisphere homologue of the cyan cluster and included large peak regions in right superior and middle frontal gyri extending into right anterior cingulate and insula; right inferior parietal lobe extending into right superior temporal gyrus; right cingulate gyrus; and right precuneus. Additional peak regions in the right hemisphere included right middle temporal gyrus; right thalamus; and right claustrum. Two peak regions were found in the left hemisphere in left inferior and superior parietal lobes and left middle and superior frontal gyri.

We further examined which network labels contributed most to which cluster, finding that SPMs labeled FPN contributed to all four clusters (Fig. 4b), SPMs labeled ECN contributed most to the green cluster, and SPMs labeled DAN and TPN contributed to the red cluster. Care must, however, be taken in interpreting these results regarding which network labels contributed most to which of the four clusters, as 118 of the 166 SPMs were labeled FPN, ECN, or DAN, meaning that SPMs corresponding to these three network labels were over-represented in the clustering analysis. The over-representation of the network labels FPN, ECN, and DAN notwithstanding, we were unable to assign a unique, existing network label to these four clusters based on these results alone.

### Network Label Recommendations

Although the above quantitative meta-analyses did not yield a clear set of recommended labels, we were able to topographically identify four distinct spatial networks. Given the current state of inconsistent labeling in the literature and the need for community consensus across studies, we propose a set of label recommendations be adopted in future studies. These recommendations include a primary label based on anatomical terminology, as endorsed by recent guidelines developed by Uddin, Yeo, and Spreng (2019), as well as a secondary label inclusive of functional descriptors. First, we recommend that the red cluster have a primary label of “Dorsal Frontoparietal Network (D-FPN)”, in agreement with Uddin et al. (2019), and a secondary label of “Dorsal Attention Network”. These labels have historically been used to refer to an executive function network consisting of bilateral frontal eye fields and intraparietal sulci (e.g., Corbetta and Shulman, 2002; Vossel et al., 2014; Yeo et al., 2011), consistent with our red cluster. We recommend that the cyan and purple clusters have a primary label of “Left Lateral Frontoparietal Network (Left L-FPN)” and “Right Lateral Frontoparietal Network (Right L-FPN)”, respectively, in agreement with Uddin et al. (2019), and a secondary label of “Left Central Executive Network” and “Right Central Executive Network”. Both sets of labels have been previously used by numerous studies to describe visually similar bilateral and unilateral networks comprised of lateral prefrontal cortical and inferior parietal regions (e.g., Chen et al., 2013; Goulden et al., 2014; Allen et al., 2011; Smith et al., 2009). Finally, we recommend a primary label of “Dorsomedial Frontoparietal Network (dM-FPN)” for the green cluster. We first considered “Medial Frontoparietal Network”, in line with terminology from Uddin et al. (2019); however, we note that this is not the same network as the canonical default mode network (Raichle, 2015). The current network emphasizes anterior and midcingulate, rather than ventromedial prefrontal cortex, and medial superior parietal, rather than posterior cingulate and precuneus. Given this, the “dorso” descriptor was added to differentiate this frontoparietal pattern from that of the default mode network. In addition, we recommend a secondary label of “Anterior Control Network”, which reflects that this cluster is comprised primarily of regions included in the anterior aspects of the Frontoparietal Control Network from the 7-Network parcellation described by Yeo et al., (Yeo et al. 2011). Further, this green cluster is also visually similar to Network 7 from the Yeo et al., 17-Network parcellation (Yeo et al. 2011), created using their dorsal anterior prefrontal cortex seed (PFCda). While the label “Anterior Control Network” is not commonly used, it is not without precedence (Langenecker et al. 2004; Khasawinah et al. 2017). Other authors (e.g., Smith et al., 2009) have previously labeled visually similar networks “Executive Control Network”, however, we intentionally did not want to re-use terms (e.g., “Executive”) across multiple different clusters.

Thresholded versions of these clustering-derived maps, suitable for use as templates in future studies, as well as unthresholded statistical maps, are available for download on NeuroVault (https://identifiers.org/neurovault.collection:8448).

## Discussion

Using an image-based meta-analytic approach, we demonstrated that the number of network labels related to executive function in the literature exceeds the number of underlying functional networks ascribed to these labels. Four topographically distinct networks were extracted from images with nine different network labels related to executive function reported across 69 different published papers. Importantly, these four networks were not consistently labeled by any of the nine possible network labels. In fact, there was very little topographical consistency within each network label. Labels such as EN, FPN, CEN, and WMN were frequently used to describe topographically similar networks. The identification of four different networks, which are frequently referred to with interchangeable and varying network labels and map to similar terms is concerning. Such inconsistencies and ambiguities may impart limitations on how we as a community of scientists discuss and interpret results in the literature and build on prior work.

While we were unable to disentangle the link between commonly used network labels and commonly reported neural networks in the literature, we were able to identify four distinct clusters from published SPMs of executive functions using image-based meta-analysis. The four clusters yielded networks that are visually similar to existing templates from Smith et al. (Smith et al. 2009) and Yeo et al. (Yeo et al. 2011) (Fig. 1), suggesting that most studies examining executive function are identifying a similar set of brain networks. However, even here, our clusters do not fully overlap with these previously published templates, potentially due to ours being derived from collating both task-based and resting-state fMRI data from publications and the use of single, large resting state fMRI datasets by Smith et al. and Yeo et al. Moving forward, we recommend that the results of our meta-analysis (accessible via NeuroVault) be used as templates in future studies to promote community consensus. We further recommend that a consistent set of labels be adopted for these four networks, including Dorsal Attention Network (DAN), Left and Right Central Executive Network (CEN), and Anterior Control Network (ACN).

Similar to other neuroimaging meta-analyses, our study is limited by concerns regarding the included studies. In particular, concern exists for the results of our first image-based meta-analysis regarding the small number of datasets for some of the network names. However, this is due to some network labels being more commonly used by authors than others. Almost seventy percent of the group-level SPMs we received were labeled as FPN, DAN, or ECN; three of the executive function network labels used by the well-cited Smith et al. (Smith et al. 2009) and Yeo et al. (Yeo et al. 2011) template sets. Our view is that full consideration of the range of network labels currently used by researchers, inclusive of even the less frequently used labels, was appropriate given our primary goal to assess variability in label usage across the field. Moreover, we note that our second image-based meta-analysis, which included clustering across SPMs regardless of assigned label, yielded four clusters reflecting four distinct spatial networks. We note that these results are well aligned with a similar set of four networks described by both Smith et al. (Smith et al. 2009) and Yeo et al. (Yeo et al. 2011). The lesser-used network labels, such as CCN, WMN, and TPN, are likely being used by researchers to describe any of these four spatial networks, leading to community challenges in effectively discussing and interpreting results across studies. Beyond variability in label assignment, we also considered that study sample size may have limited our results. To this end, we conducted ANOVA of study sample sizes to determine if there were significant differences across labels. Our analysis found no significant association between sample size and network label (*F*(148,9)=0.239; *P*=0.9881; Supplemental Fig. 6), suggesting that label assignment does not reflect prevalence of under-powered studies. However, we cannot rule out additional unknown ways in which label variability may perhaps be related to individual variability between participants. As a final limitation, 21 out of 188 studies were not able to provide their original data. Two studies, Smith et al. (Smith et al. 2009) and Allen et al. (Allen et al. 2011), had uploaded their images to an online repository; however, none of the other included studies made their data available in any publicly accessible online repository, such as NeuroVault (Gorgolewski et al. 2016). A large number of studies did not respond to our request for data sharing (83 out of 188), despite extensive efforts by the authors in following-up with at least one additional email request, retrieving current email addresses, and approaching the principal investigators when the corresponding author did not respond or had an invalid email address. This greatly hinders efforts towards reproducibility, which is a concern that is widely shared by the research community, as evidenced by the COBIDAS initiative (Nichols et al. 2016). Image-based meta-analyses offer a number of advantages over coordinate-based meta-analyses (Salimi-Khorshidi et al. 2009), yet will remain a challenge until the uploading of unthresholded study images to repositories such as NeuroVault becomes a *de facto* community standard.

## Conclusions

Our results offer insight in the inconsistent use of executive function network labels in literature. As shown through both sets of analysis, there is very little evidence indicating that a network labeled, for example, TPN by one study will have a spatial topography that is similar to another network labeled TPN by another study. Conversely, a network labeled TPN by one study may have a similar spatial topography to network labeled ECN by another study. Clustering image-based meta-analysis maps regardless of author-assigned labels identified four spatially distinct networks that are visually similar to previously published templates of executive function. Although our results indicated that there is no consistent mapping of these four networks to the network names commonly used in the literature, we describe these networks and provide labeling recommendations. To promote community consensus across studies, we further propose that researchers should adopt a labeled template set such as ours when assigning labels to executive function networks.

## Author Contributions

HvEV, STW, MCR, and ARL conceived of and designed the study. HvEV and STW performed the literature search and collected the published SPMs. TS and MCR contributed scripts, performed meta-analyses, and drafted the figures. HvEV and STW co-wrote the manuscript. All authors contributed to the revisions and approved the final version. HvEV and STW contributed equally to this work.

## Acknowledgements

The authors wish to extend a sincere thank you to all of the authors who provided their published statistical maps, thus allowing us to perform these analyses. A special thanks go to those authors who went on extended scavenger hunts through their archives in an effort to assist us with our project. ARL, MCR, and TS were supported by NSF 1631325 and NIH R01 DA041353.

