## Supplemental Table 1 for "What executive function network is that? An image-based meta-analysis of network labels"

[†]Both authors contributed equally.

**Affiliations:**

**CONTENTS:**

**Supplemental Table 1.** Full results of literature search for studies and statistical maps included in the image-based meta-analyses.

**Supplemental Figure 1.** More detailed depiction of the cluster analysis results.

**Supplemental Figure 2.** More detailed depiction of the cyan cluster (Left Central Executive Network/Left Frontoparietal Network).

**Supplemental Table 2.** Peak regions of the cyan cluster at a corrected $p < 0.05$.

**Supplemental Figure 3.** More detailed depiction of the green cluster (Anterior Control Network).

**Supplemental Table 3.** Peak regions of the green cluster at a corrected $p < 0.05$.

**Supplemental Figure 4.** More detailed depiction of the red cluster (Dorsal Attention Network).

**Supplemental Table 4.** Peak regions of the red cluster at a corrected $p < 0.05$.

**Supplemental Figure 5.** More detailed depiction of the purple cluster (Right Central Executive Network/Right Frontoparietal Network).

**Supplemental Table 5.** Peak regions of the purple cluster at a corrected $p < 0.05$.

**Supplemental Figure 6.** Results of ANOVA on study sample sizes to determine the effect of sample size for a given group-level SPM across the nine network labels of interest. The overall effect of study sample size was not significant ($F(148,9)=0.239$; $P=0.9881$).

| First Author | Publication Year | Title | Journal | DOI | PMID | Network name | Figure # | # Subjects (Healthy Controls) | # Males | Age | Task | Task Type | RDoC Construct | MNI/TAL/Other |
|---|---|---|---|---|---|---|---|---|---|---|---|---|---|---|
| Allen EA | 2011 | **A baseline for the multivariate comparison of resting-state networks** | Frontiers in systems neuroscience | 10.3389/fnsys.2011.00002 | 21442040 | FrontoParietal Network | **Figure 4 (IC 34)** | 603 | 298 | 23.4±9.2 | Rest | | | MNI |
| Allen EA | 2011 | **A baseline for the multivariate comparison of resting-state networks** | Frontiers in systems neuroscience | 10.3389/fnsys.2011.00002 | 21442040 | FrontoParietal Network | **Figure 4 (IC 60)** | 603 | 298 | 23.4±9.2 | Rest | | | MNI |
| Aso T | 2016 | **Dynamic interactions of the cortical networks during thought suppression** | Brain and Behavior | 10.1002/brb3.503 | 27547504 | Dorsal Attention Network | **Figure 1 (IC24)** | 20 | 15 | 21.5±0.9 | Task | visual imagery | Cognitive Control | MNI |
| Aso T | 2016 | **Dynamic interactions of the cortical networks during thought suppression** | Brain and Behavior | 10.1002/brb3.503 | 27547504 | Executive Control Network | **Figure 1 (IC21)** | 20 | 15 | 21.5±0.9 | Task | visual imagery | Cognitive Control | MNI |
| Aso T | 2016 | **Dynamic interactions of the cortical networks during thought suppression** | Brain and Behavior | 10.1002/brb3.503 | 27547504 | FrontoParietal Network | **Figure 1 (IC22)** | 20 | 15 | 21.5±0.9 | Task | visual imagery | Cognitive Control | MNI |
| Baggio H | 2015 | **Cognitive impairment and resting-state network connectivity in Parkinson's disease** | Human Brain Mapping | 10.1002/hbm.22622 | 25164875 | Dorsal Attention Network | **Figure 1 (Group-level Networks: HC: DAN)** | 36 | 19 | 63.4±10.5 | Rest | | | MNI |
| Baggio H | 2015 | **Cognitive impairment and resting-state network connectivity in Parkinson's disease** | Human Brain Mapping | 10.1002/hbm.22622 | 25164875 | FrontoParietal Network | **Figure 1 (Group-level Networks: HC: leftFPN)** | 36 | 19 | 63.4±10.5 | Rest | | | MNI |
| Baggio H | 2015 | **Cognitive impairment and resting-state network connectivity in Parkinson's disease** | Human Brain Mapping | 10.1002/hbm.22622 | 25164875 | FrontoParietal Network | **Figure 1 (Group-level Networks: HC: rightFPN)** | 36 | 19 | 63.4±10.5 | Rest | | | MNI |
| Bar K-J | 2016 | **Functional connectivity and network analysis of midbrain and brainstem nuclei** | NeuroImage | 10.1016/j.neuroimage.2016.03.071 | 27046112 | Executive Control Network | **Figure 4 (IC2: ECN)** | 154 | 73 | 28.8±9.6 | Rest | | | MNI |
| Barber AD | 2013 | **Developmental changes in within- and between-network connectivity between late childhood and adulthood** | Neuropsychologia | 10.1016/j.neuropsychologia.2012.11.011 | 23174403 | Task Positive Network | **Figure 1a (adults)** | 28 | 12 | 27.57±6.22 | Rest | | | MNI |
| Barnacle G | 2016 | **The list-composition effect in memory for emotional and neutral pictures: Differential contribution of ventral and dorsal attention networks to successful encoding** | Neuropsychologia | 10.1016/j.neuropsychologia.2016.06.023 | 27342257 | Dorsal Attention Network | **Figure 3** | 22 | 0 | 26 | Task | free-recall memory | Working Memory | MNI |
| Barnacle G | 2016 | **The list-composition effect in memory for emotional and neutral pictures: Differential contribution of ventral and dorsal attention networks to successful encoding** | Neuropsychologia | 10.1016/j.neuropsychologia.2016.06.023 | 27342257 | Ventral Attention Network | **Figure 2** | 22 | 0 | 26 | Task | free-recall memory | Working Memory | MNI |
| Bey K | 2015 | **Susceptibility to everyday cognitive failure is reflected in functional network interactions in the resting brain** | NeuroImage | 10.1016/j.neuroimage.2015.07.041 | 26210814 | Dorsal Attention Network | **Figure 1 (24)** | 71 | 14 | 21.56±3.28 | Rest | | | MNI |
| Bey K | 2015 | **Susceptibility to everyday cognitive failure is reflected in functional network interactions in the resting brain** | NeuroImage | 10.1016/j.neuroimage.2015.07.041 | 26210814 | Executive Control Network | **Figure 1 (15)** | 71 | 14 | 21.56±3.28 | Rest | | | MNI |
| Bey K | 2015 | **Susceptibility to everyday cognitive failure is reflected in functional network interactions in the resting brain** | NeuroImage | 10.1016/j.neuroimage.2015.07.041 | 26210814 | FrontoParietal Network | **Figure 1 (1)** | 71 | 14 | 21.56±3.28 | Rest | | | MNI |
| Bey K | 2015 | **Susceptibility to everyday cognitive failure is reflected in functional network interactions in the resting brain** | NeuroImage | 10.1016/j.neuroimage.2015.07.041 | 26210814 | FrontoParietal Network | **Figure 1 (6)** | 71 | 14 | 21.56±3.28 | Rest | | | MNI |
| Blautzik J | 2013 | **Classifying fMRI-derived resting-state connectivity patterns according to their daily rhythmicity** | NeuroImage | 10.1016/j.neuroimage.2012.08.010 | 22906784 | Dorsal Attention Network | **Figure 2 (IC 7)** | 15 | 6 | 23±3 | Rest | | | MNI |
| Blautzik J | 2013 | **Classifying fMRI-derived resting-state connectivity patterns according to their daily rhythmicity** | NeuroImage | 10.1016/j.neuroimage.2012.08.010 | 22906784 | Executive Control Network | **Figure 2 (IC 6)** | 15 | 6 | 23±3 | Rest | | | MNI |
| Blautzik J | 2013 | **Classifying fMRI-derived resting-state connectivity patterns according to their daily rhythmicity** | NeuroImage | 10.1016/j.neuroimage.2012.08.010 | 22906784 | FrontoParietal Network | **Figure 2 (IC 8)** | 15 | 6 | 23±3 | Rest | | | MNI |
| Blautzik J | 2013 | **Classifying fMRI-derived resting-state connectivity patterns according to their daily rhythmicity** | NeuroImage | 10.1016/j.neuroimage.2012.08.010 | 22906784 | FrontoParietal Network | **Figure 2 (IC 19)** | 15 | 6 | 23±3 | Rest | | | MNI |
| Braga R | 2013 | **Separable networks for top-down attention to auditory non-spatial and visuospatial modalities** | NeuroImage | 10.1016/j.neuroimage.2013.02.023 | 23435206 | FrontoParietal Network | **Figure 2 Whole Brain Auditory (third from top)** | 20 | 11 | 29.7 | Task | auditory attention | Auditory Perception | MNI |
| Braga R | 2013 | **Separable networks for top-down attention to auditory non-spatial and visuospatial modalities** | NeuroImage | 10.1016/j.neuroimage.2013.02.023 | 23435206 | FrontoParietal Network | **Figure 2 Whole Brain Visual (third from top)** | 20 | 11 | 29.7 | Task | auditory attention | Auditory Perception | MNI |
| Braunlich K | 2015 | **Frontoparietal networks involved in categorization and item working memory** | NeuroImage | 10.1016/j.neuroimage.2014.11.051 | 25482265 | Central Executive Network | **Figure 5** | 17 | 6 | 27 | Task | delayed matched to category | Working Memory | MNI |
| Bray S | 2015 | **Spatial and temporal functional connectivity changes between resting and attentive states** | Human Brain Mapping | 10.1002/hbm.22646 | 25271132 | Dorsal Attention Network | **Figure 1 (Rest (DAN))** | 16 | 7 | 22 | Rest | | | MNI |
| Bray S | 2015 | **Spatial and temporal functional connectivity changes between resting and attentive states** | Human Brain Mapping | 10.1002/hbm.22646 | 25271132 | Ventral Attention Network | **Figure 1 (Rest (VAN))** | 16 | 7 | 22 | Rest | | | MNI |
| Bray S | 2015 | **Spatial and temporal functional connectivity changes between resting and attentive states** | Human Brain Mapping | 10.1002/hbm.22646 | 25271132 | Dorsal Attention Network | **Figure 1 (Attention (DAN))** | 16 | 7 | 22 | Task | continuous visual attention | Attention | MNI |
| Bray S | 2015 | **Spatial and temporal functional connectivity changes between resting and attentive states** | Human Brain Mapping | 10.1002/hbm.22646 | 25271132 | Ventral Attention Network | **Figure 1 (Attention (VAN))** | 16 | 7 | 22 | Task | continuous visual attention | Attention | MNI |
| Cassidy CM | 2016 | **Dynamic Connectivity between Brain Networks Supports Working Memory: Relationships to Dopamine Release and Schizophrenia** | J Neuroscience | 10.1523/JNEUROSCI.3296-15.2016 | 27076432 | FrontoParietal Network | **Figure 2 (lFPN)** | 39 | 18 | 33.5±8.2 | Task | picture n-back | Working Memory | MNI |
| Cassidy CM | 2016 | **Dynamic Connectivity between Brain Networks Supports Working Memory: Relationships to Dopamine Release and Schizophrenia** | J Neuroscience | 10.1523/JNEUROSCI.3296-15.2016 | 27076432 | FrontoParietal Network | **Figure 2 (rFPN)** | 39 | 18 | 33.5±8.2 | Task | picture n-back | Working Memory | MNI |
| Costumero V | 2015 | **A new window to understanding individual differences in reward sensitivity from attentional networks** | Brain structure & function | 10.1007/s00429-014-0760-6 | 24696182 | FrontoParietal Network | **Figure 3 C09 (Experiment 1)** | 41 | 41 | 23.3±4.1 | Task | monetary incentive delay | Reward Anticipation | MNI |
| Costumero V | 2015 | **A new window to understanding individual differences in reward sensitivity from attentional networks** | Brain structure & function | 10.1007/s00429-014-0760-6 | 24696182 | FrontoParietal Network | **Figure 3 C07 (Experiment 2)** | 30 | 30 | 23.7±3 | Task | visual incentive delay | Reward Anticipation | MNI |
| Costumero V | 2015 | **How bilingualism shapes the functional architecture of the brain: A study on executive control in early bilinguals and monolinguals** | Human Brain Mapping | 10.1002/hbm.22996 | 26376449 | FrontoParietal Network | **Figure 1 C12** | 20,19 | 8,9 | 21.1±1.4, 20.5±2.9 | Task | go/no-go | Cognitive Control | MNI |
| Costumero V | 2015 | **How bilingualism shapes the functional architecture of the brain: A study on executive control in early bilinguals and monolinguals** | Human Brain Mapping | 10.1002/hbm.22996 | 26376449 | FrontoParietal Network | **Figure 1 C18** | 20,19 | 8,9 | 21.1±1.4, 20.5±2.9 | Task | go/no-go | Cognitive Control | MNI |
| De Bondt T | 2015 | **Stability of resting state networks in the female brain during hormonal changes and their relation to premenstrual symptoms** | Brain Research | 10.1016/j.brainres.2015.07.045 | 26253822 | Executive Control Network | **Figure 4** | 37 | 0 | 23.9 | Rest | | | MNI |
| De Chastelaine M | 2014 | **The relationship between task-related and subsequent memory effects** | Human Brain Mapping | 10.1002/hbm.22430 | 24615858 | Task Positive Network | **Figure 2** | 18 | 7 | 18-24 | Task | associative encoding | Working Memory | MNI |
| Deng Z | 2016 | **Resting-state low-frequency fluctuations reflect individual differences in spoken language learning** | Cortex | 10.1016/j.cortex.2015.11.020 | 26866283 | Dorsal Attention Network | **Figure 5 (DAN)** | 20 | 9 | 25.9±4.79 | Rest | | | MNI |
| Deng Z | 2016 | **Resting-state low-frequency fluctuations reflect individual differences in spoken language learning** | Cortex | 10.1016/j.cortex.2015.11.020 | 26866283 | FrontoParietal Network | **Figure 5 (FPN)** | 20 | 9 | 25.9±4.79 | Rest | | | MNI |
| Di X | 2015 | **Dynamic brain functional connectivity modulated by resting-state networks** | Brain structure & function | 10.1007/s00429-013-0634-3 | 25713839 | Executive Network | **Figure 2 (R Executive)** | 63 | 28 | 21.3 | Rest | | | MNI |
| Di X | 2015 | **Dynamic brain functional connectivity modulated by resting-state networks** | Brain structure & function | 10.1007/s00429-013-0634-3 | 25713839 | Executive Network | **Figure 2 (L Executive)** | 63 | 28 | 21.3 | Rest | | | MNI |
| Di X | 2014 | **Modulatory interactions between the default mode network and task positive networks in resting-state** | PeerJ | 10.7717/peerj.367 | 24860698 | Dorsal Attention Network | **Figure 1D** | 64 | 40 | 21.1 | Rest | | | MNI |
| Di X | 2014 | **Modulatory interactions between the default mode network and task positive networks in resting-state** | PeerJ | 10.7717/peerj.367 | 24860698 | Executive Network | **Figure 1E** | 64 | 40 | 21.1 | Rest | | | MNI |
| Di X | 2014 | **Modulatory interactions between the default mode network and task positive networks in resting-state** | PeerJ | 10.7717/peerj.367 | 24860698 | Executive Network | **Figure 1F** | 64 | 40 | 21.1 | Rest | | | MNI |
| Domagalik A | 2012 | **Neural networks related to pro-saccades and anti-saccades revealed by independent component analysis** | NeuroImage | 10.1016/j.neuroimage.2012.06.006 | 22705376 | Executive Control Network | **Figure 2 (ECN) (ProSaccadic task)** | 15 | 15 | 27.4±5.6 | Task | pro-saccade | Cognitive Control | TAL |
| Domagalik A | 2012 | **Neural networks related to pro-saccades and anti-saccades revealed by independent component analysis** | NeuroImage | 10.1016/j.neuroimage.2012.06.006 | 22705376 | FrontoParietal Network | **Figure 2 (frontoparietal right) (ProSaccadic task)** | 15 | 15 | 27.4±5.6 | Task | pro-saccade | Cognitive Control | TAL |
| Domagalik A | 2012 | **Neural networks related to pro-saccades and anti-saccades revealed by independent component analysis** | NeuroImage | 10.1016/j.neuroimage.2012.06.006 | 22705376 | FrontoParietal Network | **Figure 2 (frontoparietal left) (ProSaccadic task)** | 15 | 15 | 27.4±5.6 | Task | pro-saccade | Cognitive Control | TAL |
| Domagalik A | 2012 | **Neural networks related to pro-saccades and anti-saccades revealed by independent component analysis** | NeuroImage | 10.1016/j.neuroimage.2012.06.006 | 22705376 | Executive Control Network | **Figure 2 (ECN) (AntiSaccadic task)** | 15 | 15 | 27.4±5.6 | Task | pro-saccade | Cognitive Control | TAL |
| Domagalik A | 2012 | **Neural networks related to pro-saccades and anti-saccades revealed by independent component analysis** | NeuroImage | 10.1016/j.neuroimage.2012.06.006 | 22705376 | FrontoParietal Network | **Figure 2 (frontoparietal right) (AntiSaccadic task)** | 15 | 15 | 27.4±5.6 | Task | pro-saccade | Cognitive Control | TAL |
| Domagalik A | 2012 | **Neural networks related to pro-saccades and anti-saccades revealed by independent component analysis** | NeuroImage | 10.1016/j.neuroimage.2012.06.006 | 22705376 | FrontoParietal Network | **Figure 2 (frontoparietal left) (AntiSaccadic task)** | 15 | 15 | 27.4±5.6 | Task | pro-saccade | Cognitive Control | TAL |
| Elton A | 2015 | **Task-positive Functional Connectivity of the Default Mode Network Transcends Task Domain** | Journal of Cognitive Neuroscience | 10.1162/jocn_a_00859 | 26244722 | Dorsal Attention Network | **Figure 5A, DA** | 20 | 9 | 28.5 | Rest | | | MNI |
| Elton A | 2015 | **Task-positive Functional Connectivity of the Default Mode Network Transcends Task Domain** | Journal of Cognitive Neuroscience | 10.1162/jocn_a_00859 | 26244722 | FrontoParietal Network | **Figure 5A, RF** | 20 | 9 | 28.5 | Rest | | | MNI |
| Elton A | 2015 | **Task-positive Functional Connectivity of the Default Mode Network Transcends Task Domain** | Journal of Cognitive Neuroscience | 10.1162/jocn_a_00859 | 26244722 | FrontoParietal Network | **Figure 5A, LF** | 20 | 9 | 28.5 | Rest | | | MNI |
| Evers EA | 2012 | **The Effects of Sustained Cognitive Task Performance on Subsequent Resting State Functional Connectivity in Healthy Young and Middle-Aged Male Schoolteachers** | Brain Connectivity | 10.1089/brain.2011.0060 | 22497341 | Executive Control Network | **Figure 2A** | 32 | 32 | | Rest | | | MNI |
| Evers EA | 2012 | **The Effects of Sustained Cognitive Task Performance on Subsequent Resting State Functional Connectivity in Healthy Young and Middle-Aged Male Schoolteachers** | Brain Connectivity | 10.1089/brain.2011.0060 | 22497341 | FrontoParietal Network | **Figure 2B** | 32 | 32 | | Rest | | | MNI |
| Evers EA | 2012 | **The Effects of Sustained Cognitive Task Performance on Subsequent Resting State Functional Connectivity in Healthy Young and Middle-Aged Male Schoolteachers** | Brain Connectivity | 10.1089/brain.2011.0060 | 22497341 | FrontoParietal Network | **Figure 2C** | 32 | 32 | | Rest | | | MNI |
| Filippi M | 2013 | **The organization of intrinsic brain activity differs between genders: A resting-state fMRI study in a large cohort of young healthy subjects** | Human Brain Mapping | 10.1002/hbm.21514 | 22359372 | Executive Control Network | **Figure 2G** | 104 | 48 | 20-29 | Rest | | | MNI |
| Filippi M | 2013 | **The organization of intrinsic brain activity differs between genders: A resting-state fMRI study in a large cohort of young healthy subjects** | Human Brain Mapping | 10.1002/hbm.21514 | 22359372 | Working Memory Network | **Figure 2K** | 104 | 48 | 20-29 | Rest | | | MNI |
| Filippi M | 2013 | **The organization of intrinsic brain activity differs between genders: A resting-state fMRI study in a large cohort of young healthy subjects** | Human Brain Mapping | 10.1002/hbm.21514 | 22359372 | Working Memory Network | **Figure 2L** | 104 | 48 | 20-29 | Rest | | | MNI |
| Franco A | 2013 | **Impact of analysis methods on the reproducibility and reliability of resting-state networks** | Brain Connectivity | 10.1089/brain.2012.0134 | 23705789 | Executive Control Network | **Figure 3 (Executive Control: Model)** | 30,19 | 21,10 | 31.83±10.72,28.95±10.02 | Rest | | | TAL |
| Franco A | 2013 | **Impact of analysis methods on the reproducibility and reliability of resting-state networks** | Brain Connectivity | 10.1089/brain.2012.0134 | 23705789 | Executive Control Network | **Figure 3 (Executive Control: Testing)** | 30,19 | 21,10 | 31.83±10.72,28.95±10.02 | Rest | | | TAL |
| Franco A | 2013 | **Impact of analysis methods on the reproducibility and reliability of resting-state networks** | Brain Connectivity | 10.1089/brain.2012.0134 | 23705789 | FrontoParietal Network | **Figure 3 (Right Frontoparietal: Model)** | 30,19 | 21,10 | 31.83±10.72,28.95±10.02 | Rest | | | TAL |
| Franco A | 2013 | **Impact of analysis methods on the reproducibility and reliability of resting-state networks** | Brain Connectivity | 10.1089/brain.2012.0134 | 23705789 | FrontoParietal Network | **Figure 3 (Right Frontoparietal: Testing)** | 30,19 | 21,10 | 31.83±10.72,28.95±10.02 | Rest | | | TAL |
| Franco A | 2013 | **Impact of analysis methods on the reproducibility and reliability of resting-state networks** | Brain Connectivity | 10.1089/brain.2012.0134 | 23705789 | FrontoParietal Network | **Figure 3 (Left Frontoparietal: Model)** | 30,19 | 21,10 | 31.83±10.72,28.95±10.02 | Rest | | | TAL |
| Franco A | 2013 | **Impact of analysis methods on the reproducibility and reliability of resting-state networks** | Brain Connectivity | 10.1089/brain.2012.0134 | 23705789 | FrontoParietal Network | **Figure 3 (Left Frontoparietal: Testing)** | 30,19 | 21,10 | 31.83±10.72,28.95±10.02 | Rest | | | TAL |
| Fuentes-Claramonte P | 2016 | **Characterizing individual differences in reward sensitivity from the brain networks involved in response inhibition** | NeuroImage | 10.1016/j.neuroimage.2015.08.067 | 26343318 | FrontoParietal Network | **Figure 2 (IC12)** | 61 | 37 | | Task | stop-signal | Cognitive Control | MNI |
| Fuentes-Claramonte P | 2016 | **Characterizing individual differences in reward sensitivity from the brain networks involved in response inhibition** | NeuroImage | 10.1016/j.neuroimage.2015.08.067 | 26343318 | FrontoParietal Network | **Figure 2 (IC25)** | 61 | 37 | | Task | stop-signal | Cognitive Control | MNI |
| Goya-Maldonado R | 2016 | **Differentiating unipolar and bipolar depression by alterations in large-scale brain networks** | Human Brain Mapping | 10.1002/hbm.23070 | 26611711 | FrontoParietal Network | **Figure 2 (FP, Control- yellow)** | 20 | 7 | 36.2±11.3 | Rest | | | MNI |
| Harding IH | 2016 | **Cortical Representations of Cognitive Control and Working Memory Are Dependent Yet Non-Interacting** | Cerebral Cortex | 10.1093/cercor/bhu208 | 25249406 | Cognitive Control Network | **Figure 3, S1** | 25 | 14 | 25.5±4.4 | Task | n-back/distractor | Working Memory/Cognitive Control | MNI |
| Harding IH | 2016 | **Cortical Representations of Cognitive Control and Working Memory Are Dependent Yet Non-Interacting** | Cerebral Cortex | 10.1093/cercor/bhu208 | 25249406 | Working Memory Network | **Figure 3, S1** | 25 | 14 | 25.5±4.4 | Task | n-back/distractor | Working Memory/Cognitive Control | MNI |
| Harding IH | 2016 | **Fronto-Cerebellar Dysfunction and Dysconnectivity Underlying Cognition in Friedreich Ataxia: The IMAGE-FRDA Study** | Human Brain Mapping | 10.1002/hbm.23034 | 26502936 | Cognitive Control Network | **Figure S3, CON** | 34 | 17 | 33.6 | Task | n-back | Working Memory | MNI |
| Heinzel S | 2014 | **Working Memory Load-Dependent Brain Response Predicts Behavioral Training Gains in Older Adults** | J Neuroscience | 10.1523/JNEUROSCI.2463-13.2014 | 24453314 | Working Memory Network | **Figure 1B** | 18,19 | 10,13 | 24.1±2.43, 66±3.73 | Task | n-back | Working Memory | MNI |
| Hiltunen T | 2014 | **Infra-slow EEG fluctuations are correlated with resting-state network dynamics in fMRI** | J Neuroscience | 10.1523/JNEUROSCI.0276-13.2014 | 24403137 | Dorsal Attention Network | **Figure 2 (red-yellow, DAN)** | 21 | 14 | 24.1±2.8 | Rest | | | MNI |
| Hiltunen T | 2014 | **Infra-slow EEG fluctuations are correlated with resting-state network dynamics in fMRI** | J Neuroscience | 10.1523/JNEUROSCI.0276-13.2014 | 24403137 | Executive Network | **Figure 2 (red-yellow, EXEC)** | 21 | 14 | 24.1±2.8 | Rest | | | MNI |
| Huang H | 2015 | **Linking functional connectivity and structural connectivity quantitatively: A comparison of methods** | Brain Connectivity | 10.1089/brain.2015.0382 | 26598788 | Central Executive Network | **Figure 3A (Left Central Executive Network)** | 12 | 7 | 25.4±2.5 | Rest | | | MNI |
| Huang H | 2015 | **Linking functional connectivity and structural connectivity quantitatively: A comparison of methods** | Brain Connectivity | 10.1089/brain.2015.0382 | 26598788 | Central Executive Network | **Figure 3A (Right Central Executive Network)** | 12 | 7 | 25.4±2.5 | Rest | | | MNI |
| Jolles D | 2011 | **A comprehensive study of whole-brain functional connectivity in children and young adults** | Cerebral Cortex | 10.1093/cercor/bhq104 | 20542991 | Dorsal Attention Network | **Figure 1G, purple (adults)** | 29 | 13 | 22.2±1.67 | Rest | | | MNI |
| Jolles D | 2011 | **A comprehensive study of whole-brain functional connectivity in children and young adults** | Cerebral Cortex | 10.1093/cercor/bhq104 | 20542991 | Executive Control Network | **Figure 1F, purple (adults)** | 29 | 13 | 22.2±1.67 | Rest | | | MNI |
| Jolles D | 2011 | **A comprehensive study of whole-brain functional connectivity in children and young adults** | Cerebral Cortex | 10.1093/cercor/bhq104 | 20542991 | FrontoParietal Network | **Figure 1H, purple (adults)** | 29 | 13 | 22.2±1.67 | Rest | | | MNI |
| Jolles D | 2011 | **A comprehensive study of whole-brain functional connectivity in children and young adults** | Cerebral Cortex | 10.1093/cercor/bhq104 | 20542991 | FrontoParietal Network | **Figure 1I, purple (adults)** | 29 | 13 | 22.2±1.67 | Rest | | | MNI |
| Karahanoglu F | 2015 | **Transient brain activity disentangles fMRI resting-state dynamics in terms of spatially and temporally overlapping networks** | Nature Communication | 10.1038/ncomms8751 | 26178017 | Dorsal Attention Network | **Figure 2a (iCAP 6)** | 14 | | 38.4±6 | Rest | | | MNI |
| Karahanoglu F | 2015 | **Transient brain activity disentangles fMRI resting-state dynamics in terms of spatially and temporally overlapping networks** | Nature Communication | 10.1038/ncomms8751 | 26178017 | Executive Network | **Figure 2a (iCAP 9)** | 14 | | 38.4±6 | Rest | | | MNI |
| Karahanoglu F | 2015 | **Transient brain activity disentangles fMRI resting-state dynamics in terms of spatially and temporally overlapping networks** | Nature Communication | 10.1038/ncomms8751 | 26178017 | FrontoParietal Network | **Figure 2a (iCAP 2)** | 14 | | 38.4±6 | Rest | | | MNI |
| Kelly AC | 2008 | **Competition between functional brain networks mediates behavioral variability** | NeuroImage | 10.1016/j.neuroimage.2007.08.008 | 17919929 | Task Positive Network | **Figure 3b (activation)** | 26 | | 28.1±8.5 | Task | flanker task | Cognitive Control | MNI |
| Kepinska O | 2009 | **Whole-brain functional connectivity during acquisition of novel grammar** | Behavioural Brain Research | 10.1016/j.bbr.2016.12.015 | 1664328 | Task Positive Network | **Figure 2a** | 42 | 10 | 23 | Rest | | | MNI |
| Kepinska O | 2009 | **Whole-brain functional connectivity during acquisition of novel grammar** | Behavioural Brain Research | 10.1016/j.bbr.2016.12.015 | 1664328 | Working Memory Network | **Figure 2c** | 42 | 10 | 23 | Rest | | | MNI |
| Koch G | 2012 | **fMRI Resting Slow Fluctuations Correlate with the Activity of Fast Cortico-Cortical Physiological Connections** | PlosOne | 10.1371/journal.pone.0052660 | 23285136 | Dorsal Attention Network | **Figure 2 (DAN)** | 19 | 7 | 21-36 | Rest | | | MNI |
| Koch G | 2012 | **fMRI Resting Slow Fluctuations Correlate with the Activity of Fast Cortico-Cortical Physiological Connections** | PlosOne | 10.1371/journal.pone.0052660 | 23285136 | FrontoParietal Network | **Figure 2 (LFPN)** | 19 | 7 | 21-36 | Rest | | | MNI |
| Koch G | 2012 | **fMRI Resting Slow Fluctuations Correlate with the Activity of Fast Cortico-Cortical Physiological Connections** | PlosOne | 10.1371/journal.pone.0052660 | 23285136 | FrontoParietal Network | **Figure 2 (RFPN)** | 19 | 7 | 21-36 | Rest | | | MNI |
| Kragel JE | 2015 | **Functional interactions between large-scale networks during memory search** | Cerebral Cortex | 10.1093/cercor/bht258 | 24084128 | Dorsal Attention Network | **Figure 3, DFP/IC7** | 20 | 8 | 18-35 | Task | free-recall memory | Working Memory | MNI |
| Kragel JE | 2015 | **Functional interactions between large-scale networks during memory search** | Cerebral Cortex | 10.1093/cercor/bht258 | 24084128 | FrontoParietal Network | **Figure 2 IC14** | 20 | 8 | 18-35 | Task | free-recall memory | Working Memory | MNI |
| Krug M | 2010 | **Adding fear to conflict: a general purpose cognitive control network is modulated by trait anxiety** | Cognitive, affective & behavioral neuroscience | 10.3758/CABN.10.3.357 | 20805537 | Cognitive Control Network | **Figure 5 (I-C nonemotional task)** | 30 | 15 | 25.4±4.7 | Task | stroop task | Cognitive Control | MNI |
| Krug M | 2010 | **Adding fear to conflict: a general purpose cognitive control network is modulated by trait anxiety** | Cognitive, affective & behavioral neuroscience | 10.3758/CABN.10.3.357 | 20805537 | Cognitive Control Network | **Figure 5 (I-C emotional task)** | 30 | 15 | 25.4±4.7 | Task | stroop task | Cognitive Control | MNI |
| Krug M | 2012 | **Proactive and reactive control during emotional interference and its relationship to trait anxiety** | Brain Research | 10.1016/j.brainres.2012.08.045 | 22960116 | Cognitive Control Network | **Figure 4 (I-C, LE task; 4a)** | 42 | 16 | 23.2±4.6 | Task | stroop task | Cognitive Control | MNI |
| Liu Y | 2016 | **Top-down Modulation of Neural Activity in Anticipatory Visual Attention: Control Mechanisms Revealed by Simultaneous EEG-fMRI** | Cerebral Cortex | 10.1093/cercor/bhu204 | 25205663 | Dorsal Attention Network | **Figure 2D** | 13 | 8 | 19±1.34 | Task | visual-spatial attention | Attention | MNI |
| Luo Y | 2014 | **Emotion perception and executive control interact in the salience network during emotionally charged working memory processing** | Human Brain Mapping | 10.1002/hbm.22573 | 25044711 | FrontoParietal Network | **Figure 3A, fronto-parietal network (2- vs 0-back)** | 25 | | 23.14±1.83 | Task | n-back | Working Memory | MNI |
| Maki-Marttunen V | 2014 | **Low attentional engagement makes attention network activity susceptible to emotional interference** | Neuroreport | 10.1097/WNR.0000000000000223 | 24999907 | Dorsal Attention Network | **Figure 1a (Main effect of attention load), Dorsal frontoparietal attention network** | 14 | 5 | 32 | Task | visual-spatial attention | Attention | MNI |
| Martinez K | 2013 | **Changes in resting-state functionally connected parietofrontal networks after videogame practice** | Human Brain Mapping | 10.1002/hbm.22129 | 22807280 | FrontoParietal Network | **Figure 2 (IC1)** | 20 | 0 | 18.95±2.65 | Rest | | | MNI |
| Martinez K | 2013 | **Changes in resting-state functionally connected parietofrontal networks after videogame practice** | Human Brain Mapping | 10.1002/hbm.22129 | 22807280 | FrontoParietal Network | **Figure 2 (IC21)** | 20 | 0 | 18.95±2.65 | Rest | | | MNI |
| Martinez K | 2013 | **Changes in resting-state functionally connected parietofrontal networks after videogame practice** | Human Brain Mapping | 10.1002/hbm.22129 | 22807280 | FrontoParietal Network | **Figure 2 (IC24)** | 20 | 0 | 18.95±2.65 | Rest | | | MNI |
| Martinez K | 2013 | **Changes in resting-state functionally connected parietofrontal networks after videogame practice** | Human Brain Mapping | 10.1002/hbm.22129 | 22807280 | FrontoParietal Network | **Figure 2 (IC25)** | 20 | 0 | 18.95±2.65 | Rest | | | MNI |
| Park B | 2012 | **Are brain networks stable during a 24-hour period?** | NeuroImage | 10.1016/j.neuroimage.2011.07.049 | 21807101 | FrontoParietal Network | **Figure 2C** | 12 | 9 | 25.42±2.84 | Rest | | | MNI |
| Park B | 2012 | **Are brain networks stable during a 24-hour period?** | NeuroImage | 10.1016/j.neuroimage.2011.07.049 | 21807101 | FrontoParietal Network | **Figure 2D** | 12 | 9 | 25.42±2.84 | Rest | | | MNI |
| Petersen N | 2014 | **Oral contraceptive pill use and menstrual cycle phase are associated with altered resting state functional connectivity** | NeuroImage | 10.1016/j.neuroimage.2013.12.016 | 24365676 | Executive Control Network | **Figure 2** | 91 | 0 | 18-40 | Rest | | | MNI |
| Reineberg A | 2015 | **Resting-state networks predict individual differences in common and specific aspects of executive function** | NeuroImage | 10.1016/j.neuroimage.2014.09.045 | 25281800 | FrontoParietal Network | **Figure 2 (FPN: IC5)** | 91 | 43 | 22.3±9.92 | Rest | | | MNI |
| Reineberg A | 2015 | **Resting-state networks predict individual differences in common and specific aspects of executive function** | NeuroImage | 10.1016/j.neuroimage.2014.09.045 | 25281800 | FrontoParietal Network | **Figure 2 (FPN: IC9)** | 91 | 43 | 22.3±9.92 | Rest | | | MNI |
| Reineberg A | 2015 | **Resting-state networks predict individual differences in common and specific aspects of executive function** | NeuroImage | 10.1016/j.neuroimage.2014.09.045 | 25281800 | Dorsal Attention Network | **Figure 2 (DAN: IC14)** | 91 | 43 | 22.3±9.92 | Rest | | | MNI |
| Reineberg A | 2015 | **Resting-state networks predict individual differences in common and specific aspects of executive function** | NeuroImage | 10.1016/j.neuroimage.2014.09.045 | 25281800 | Dorsal Attention Network | **Figure 2 (DAN: IC17)** | 91 | 43 | 22.3±9.92 | Rest | | | MNI |
| Reineberg A | 2015 | **Resting-state networks predict individual differences in common and specific aspects of executive function** | NeuroImage | 10.1016/j.neuroimage.2014.09.045 | 25281800 | Dorsal Attention Network | **Figure 2 (DAN: IC26)** | 91 | 43 | 22.3±9.92 | Rest | | | MNI |
| Reineberg A | 2015 | **Resting-state networks predict individual differences in common and specific aspects of executive function** | NeuroImage | 10.1016/j.neuroimage.2014.09.045 | 25281800 | Ventral Attention Network | **Figure 2 (VAN: IC4)** | 91 | 43 | 22.3±9.92 | Rest | | | MNI |
| Rosazza C | 2012 | **Functional connectivity during resting-state functional MR imaging: study of the correspondence between independent component analysis and region-of-interest-based methods** | AJNR. American journal of neuroradiology | 10.3174/ajnr.A2733 | 21998099 | Executive Control Network | **Figure 2 (Component 19)** | 40 | 19 | 40.8±9.3 | Rest | | | MNI |
| Rosazza C | 2012 | **Functional connectivity during resting-state functional MR imaging: study of the correspondence between independent component analysis and region-of-interest-based methods** | AJNR. American journal of neuroradiology | 10.3174/ajnr.A2733 | 21998099 | FrontoParietal Network | **Figure 2 (Component 2)** | 40 | 19 | 40.8±9.3 | Rest | | | MNI |
| Rosazza C | 2012 | **Functional connectivity during resting-state functional MR imaging: study of the correspondence between independent component analysis and region-of-interest-based methods** | AJNR. American journal of neuroradiology | 10.3174/ajnr.A2733 | 21998099 | FrontoParietal Network | **Figure 2 (Component 3)** | 40 | 19 | 40.8±9.3 | Rest | | | MNI |
| Sala-Llonch R | 2012 | **Brain connectivity during resting state and subsequent working memory task predicts behavioural performance** | Cortex | 10.1016/j.cortex.2011.07.006 | 21872853 | Working Memory Network | **Figure 4A (IC1)** | 16 | 7 | 21.31±2.41 | Task | n-back | Working Memory | MNI |
| Santangelo V | 2015 | **Parietal cortex integrates contextual and saliency signals during the encoding of natural scenes in working memory** | Human Brain Mapping | 10.1002/hbm.22984 | 26333392 | FrontoParietal Network | **Figure 2 (red)** | 14 | 4 | 22 | Task | spatial working memory | Working Memory | MNI |
| Santangelo V | 2015 | **Parietal cortex integrates contextual and saliency signals during the encoding of natural scenes in working memory** | Human Brain Mapping | 10.1002/hbm.22984 | 26333392 | FrontoParietal Network | **Figure 2 (blue)** | 14 | 4 | 22 | Task | spatial working memory | Working Memory | MNI |
| Savio A | 2017 | **Resting state networks as simultaneously measured with fMRI and PET** | Journal of Nuclear Medicine | 10.2967/jnumed.116.185835 | 28254868 | Central Executive Network | **Figure 1G (fMRI)** | 22 | 15 | 54.5±10 | Rest | | | MNI |
| Savio A | 2017 | **Resting state networks as simultaneously measured with fMRI and PET** | Journal of Nuclear Medicine | 10.2967/jnumed.116.185835 | 28254868 | Central Executive Network | **Figure 1J (fMRI)** | 22 | 15 | 54.5±10 | Rest | | | MNI |
| Savio A | 2017 | **Resting state networks as simultaneously measured with fMRI and PET** | Journal of Nuclear Medicine | 10.2967/jnumed.116.185835 | 28254868 | Executive Control Network | **Figure 1I (fMRI)** | 22 | 15 | 54.5±10 | Rest | | | MNI |
| Schulz KP | 2014 | **Emotional bias of cognitive control in adults with childhood attention-deficit/hyperactivity disorder** | NeuroImage Clinical | 10.1016/j.nicl.2014.05.016 | 24918067 | Cognitive Control Network | **Figure 2, Comparison** | 14 | 14 | 22.8±2.7 | Task | go/no-go | Cognitive Control | MNI |
| Seeley W | 2007 | **Dissociable intrinsic connectivity networks for salience processing and executive control** | J Neuroscience | 10.1523/JNEUROSCI.5587-06.2007 | 17329432 | Executive Control Network | **Figure 2 (blue-green)** | 21 | 10 | 34.6±15.8 | Rest | | | MNI |
| Seminowicz D | 2011 | **Effective treatment of chronic low back pain in humans reverses abnormal brain anatomy and function** | J Neuroscience | 10.1523/JNEUROSCI.5280-10.2011 | 21593339 | Task Positive Network | **Figure 4A (Controls)** | 16 | 8 | 40±13.16 | Task | multi-source interference task | Cognitive Control | MNI |
| Shaurya Prakash R | 2013 | **Mindfulness disposition and default-mode network connectivity in older adults** | Social Cognitive and Affective Neuroscience | 10.1093/scan/nss115 | 23051900 | Executive Network | **Figure 1D** | 25 | 6 | 65.88 | Rest | | | MNI |
| Shaurya Prakash R | 2013 | **Mindfulness disposition and default-mode network connectivity in older adults** | Social Cognitive and Affective Neuroscience | 10.1093/scan/nss115 | 23051900 | FrontoParietal Network | **Figure 1B** | 25 | 6 | 65.88 | Rest | | | MNI |
| Shaurya Prakash R | 2013 | **Mindfulness disposition and default-mode network connectivity in older adults** | Social Cognitive and Affective Neuroscience | 10.1093/scan/nss115 | 23051900 | FrontoParietal Network | **Figure 1C** | 25 | 6 | 65.88 | Rest | | | MNI |
| Smith SM | 2009 | **Correspondence of the brain's functional architecture during activation and rest** | PNAS | 10.1073/pnas.0905267106 | 19620724 | Executive Control Network | **Figure 1 (8, RSN)** | 36 | | | Rest | | | MNI |
| Smith SM | 2009 | **Correspondence of the brain's functional architecture during activation and rest** | PNAS | 10.1073/pnas.0905267106 | 19620724 | FrontoParietal Network | **Figure 1 (9, RSN)** | 36 | | | Rest | | | MNI |
| Smith SM | 2009 | **Correspondence of the brain's functional architecture during activation and rest** | PNAS | 10.1073/pnas.0905267106 | 19620724 | FrontoParietal Network | **Figure 1 (10, RSN)** | 36 | | | Rest | | | MNI |
| Spreng RN | 2012 | **Intrinsic Architecture Underlying the Relations among the Default , Dorsal Attention , and Frontoparietal Control Networks of the Human Brain** | Journal of Cognitive Neuroscience | 10.1162/jocn_a_00281 | 22905821 | Dorsal Attention Network | **Figure 1B** | 105 | 51 | 23.3±2.2 | Rest | | | MNI |
| Spreng RN | 2012 | **Intrinsic Architecture Underlying the Relations among the Default , Dorsal Attention , and Frontoparietal Control Networks of the Human Brain** | Journal of Cognitive Neuroscience | 10.1162/jocn_a_00281 | 22905821 | FrontoParietal Control Network | **Figure 1C** | 105 | 51 | 23.3±2.2 | Rest | | | MNI |
| Spreng RN | 2010 | **Default network activity, coupled with the frontoparietal control network, supports goal-directed cognition** | NeuroImage | 10.1016/j.neuroimage.2010.06.016 | 20600998 | Dorsal Attention Network | **Figure 2A (red: visuospatial planning task at TR 6)** | 20 | 3 | 21.3±3.2 | Task | visuospatial planning | Cognitive Control | MNI |
| Spreng RN | 2010 | **Default network activity, coupled with the frontoparietal control network, supports goal-directed cognition** | NeuroImage | 10.1016/j.neuroimage.2010.06.016 | 20600998 | FrontoParietal Control Network | **Figure 3A (green: both planning tasks at TR 6)** | 20 | 3 | 21.3±3.2 | Task | | | MNI |
| St. Jacques PL | 2011 | **Dynamic neural networks supporting memory retrieval** | NeuroImage | 10.1016/j.neuroimage.2011.04.039 | 21550407 | FrontoParietal Network | **Figure 1 (Frontoparietal)** | 14 | 7 | 24.43±3.73 | Task | autobiographical memory recall | Perception and Understanding of Self | MNI |
| Vacchi L | 2016 | **Working memory network dysfunction in relapse-onset multiple sclerosis phenotypes: A clinical-imaging evaluation** | Multiple Sclerosis Journal | 10.1177/1352458516656809 | 27354020 | Working Memory Network | **Figure 1a HC** | 24 | 12 | 37.6±12.2 | Task | n-back | Working Memory | MNI |
| Van Overwalle F | 2017 | **Tell me twice: A Multi-Study analysis of the functional connectivity between the cerebrum and cerebellum after repeated trait information** | NeuroImage | 10.1016/j.neuroimage.2016.08.046 | 27566262 | Executive Network | **Figure 1 (target > prime contrast of trait-implying sentences)** | 146 | | | Task | repeated trait information | Social Communication | MNI |
| Wang L | 2015 | **Large-scale functional brain network changes in taxi drivers: Evidence from resting-state fMRI** | Human Brain Mapping | 10.1002/hbm.22670 | 25338709 | FrontoParietal Network | **Figure 1C** | 40 | 37 | 40 | Rest | | | MNI |
| Wang L | 2015 | **Large-scale functional brain network changes in taxi drivers: Evidence from resting-state fMRI** | Human Brain Mapping | 10.1002/hbm.22670 | 25338709 | FrontoParietal Network | **Figure 1D** | 40 | 37 | 40 | Rest | | | MNI |
| Wisner K | 2013 | **Neurometrics of intrinsic connectivity networks at rest using fMRI: Retest reliability and cross-validation using a meta-level method** | NeuroImage | 10.1016/j.neuroimage.2013.02.066 | 23507379 | Executive Network | **Figure 4 (REF 03)** | 27 | 19 | 25±6.7 | Rest | | | MNI |
| Wisner K | 2013 | **Neurometrics of intrinsic connectivity networks at rest using fMRI: Retest reliability and cross-validation using a meta-level method** | NeuroImage | 10.1016/j.neuroimage.2013.02.066 | 23507379 | Executive Network | **Figure 4 (REF 05)** | 27 | 19 | 25±6.7 | Rest | | | MNI |
| Xu J | 2014 | **Opposite modulation of brain functional networks implicated at low vs. high demand of attention and working memory** | PlosOne | 10.1371/journal.pone.0087078 | 24498021 | Dorsal Attention Network | **Figure 2A (IC13)** | 23 | 12 | 23-41 | Rest | | | MNI |
| Xu J | 2014 | **Opposite modulation of brain functional networks implicated at low vs. high demand of attention and working memory** | PlosOne | 10.1371/journal.pone.0087078 | 24498021 | Executive Control Network | **Figure 2A (IC12)** | 23 | 12 | 23-41 | Rest | | | MNI |

| | | | | | | | | | | | | | | |
|---|---|---|---|---|---|---|---|---|---|---|---|---|---|---|
| Xu J | 2014 | **Opposite modulation of brain functional networks implicated at low vs. high demand of attention and working memory** | PlosOne | 10.1371/journal.pone.0087078 | 24498021 | FrontoParietal Network | **Figure 1A (IC4)** | 23 | 12 | 23-41 | Rest | | | MNI |
| Xu J | 2014 | **Opposite modulation of brain functional networks implicated at low vs. high demand of attention and working memory** | PlosOne | 10.1371/journal.pone.0087078 | 24498021 | FrontoParietal Network | **Figure 2A (IC22)** | 23 | 12 | 23-41 | Rest | | | MNI |
| Xu J | 2014 | **Opposite modulation of brain functional networks implicated at low vs. high demand of attention and working memory** | PlosOne | 10.1371/journal.pone.0087078 | 24498021 | FrontoParietal Network | **not depicted IC15 - RFPN** | 23 | 12 | 23-41 | Rest | | | MNI |
| Xuan B | 2016 | **The activation of interactive attentional networks** | NeuroImage | 10.1016/j.neuroimage.2016.01.017 | 26794640 | FrontoParietal Network | **Figure 5** | 24 | 13 | 26.3 | Task | revised attention network task | Attention | MNI |
| Yakunina N | 2015 | **Effects of scanner acoustic noise on intrinsic brain activity during auditory stimulation** | Neuroradiology | 10.1007/s00234-015-1561-1 | 26193957 | Dorsal Attention Network | **Figure 2 (DAN) (Sparse Temporal Acquisition)** | 14 | 8 | 30.6±4.7 | Rest | | | MNI |
| Yakunina N | 2015 | **Effects of scanner acoustic noise on intrinsic brain activity during auditory stimulation** | Neuroradiology | 10.1007/s00234-015-1561-1 | 26193957 | FrontoParietal Network | **Figure 2 (LFPN) (Sparse Temporal Acquisition)** | 14 | 8 | 30.6±4.7 | Rest | | | MNI |
| Yakunina N | 2015 | **Effects of scanner acoustic noise on intrinsic brain activity during auditory stimulation** | Neuroradiology | 10.1007/s00234-015-1561-1 | 26193957 | FrontoParietal Network | **Figure 2 (RFPN) (Sparse Temporal Acquisition)** | 14 | 8 | 30.6±4.7 | Rest | | | MNI |
| Yakunina N | 2015 | **Effects of scanner acoustic noise on intrinsic brain activity during auditory stimulation** | Neuroradiology | 10.1007/s00234-015-1561-1 | 26193957 | Ventral Attention Network | **Figure 2 (VAN) (Sparse Temporal Acquisition)** | 14 | 8 | 30.6±4.7 | Rest | | | MNI |
| Yakunina N | 2015 | **Effects of scanner acoustic noise on intrinsic brain activity during auditory stimulation** | Neuroradiology | 10.1007/s00234-015-1561-1 | 26193957 | Dorsal Attention Network | **Figure 2 (DAN) (Continuous Acquisition)** | 14 | 8 | 30.6±4.7 | Rest | | | MNI |
| Yakunina N | 2015 | **Effects of scanner acoustic noise on intrinsic brain activity during auditory stimulation** | Neuroradiology | 10.1007/s00234-015-1561-1 | 26193957 | FrontoParietal Network | **Figure 2 (LFPN) (Continuous Acquisition)** | 14 | 8 | 30.6±4.7 | Rest | | | MNI |
| Yakunina N | 2015 | **Effects of scanner acoustic noise on intrinsic brain activity during auditory stimulation** | Neuroradiology | 10.1007/s00234-015-1561-1 | 26193957 | FrontoParietal Network | **Figure 2 (RFPN) (Continuous Acquisition)** | 14 | 8 | 30.6±4.7 | Rest | | | MNI |
| Yakunina N | 2015 | **Effects of scanner acoustic noise on intrinsic brain activity during auditory stimulation** | Neuroradiology | 10.1007/s00234-015-1561-1 | 26193957 | Ventral Attention Network | **Figure 2 (VAN) (Continuous Acquisition)** | 14 | 8 | 30.6±4.7 | Rest | | | MNI |
| Zhang S | 2012 | **Functional networks for cognitive control in a stop signal task: Independent component analysis** | Human Brain Mapping | 10.1002/hbm.21197 | 21365716 | FrontoParietal Network | **Figure 2a (right FPN #4)** | 59 | 30 | 22-42 | Task | stop-signal | Cognitive Control | MNI |
| Zhang S | 2012 | **Functional networks for cognitive control in a stop signal task: Independent component analysis** | Human Brain Mapping | 10.1002/hbm.21197 | 21365716 | FrontoParietal Network | **Figure 3a (left FPN #8)** | 59 | 30 | 22-42 | Task | stop-signal | Cognitive Control | MNI |
| Zhu X | 2015 | **Model-free functional connectivity and impulsivity correlates of alcohol dependence: a resting-state study** | Addiction biology | 10.1111/adb.12272 | 26040546 | Executive Control Network | **Figure 1 (HC: LECN, green: healthy control group)** | 26 | 11 | 26.3±3.9 | Rest | | | MNI |
| Zhu X | 2015 | **Model-free functional connectivity and impulsivity correlates of alcohol dependence: a resting-state study** | Addiction biology | 10.1111/adb.12272 | 26040546 | Executive Control Network | **Figure 1 (HC: RECN, green: healthy control group)** | 26 | 11 | 26.3±3.9 | Rest | | | MNI |
| Zuo X | 2010 | **Reliable intrinsic connectivity networks: Test-retest evaluation using ICA and dual regression approach** | NeuroImage | 10.1016/j.neuroimage.2009.10.080 | 19896537 | Executive Control Network | **Figure 2 (IC08)** | 26 | 11 | 20.5±4.8 | Rest | | | MNI |
| Zuo X | 2010 | **Reliable intrinsic connectivity networks: Test-retest evaluation using ICA and dual regression approach** | NeuroImage | 10.1016/j.neuroimage.2009.10.080 | 19896537 | FrontoParietal Network | **Figure 2 (IC07)** | 26 | 11 | 20.5±4.8 | Rest | | | MNI |
| Zuo X | 2010 | **Reliable intrinsic connectivity networks: Test-retest evaluation using ICA and dual regression approach** | NeuroImage | 10.1016/j.neuroimage.2009.10.080 | 19896537 | FrontoParietal Network | **Figure 2 (IC19)** | 26 | 11 | 20.5±4.8 | Rest | | | MNI |

Supplemental Figure 1

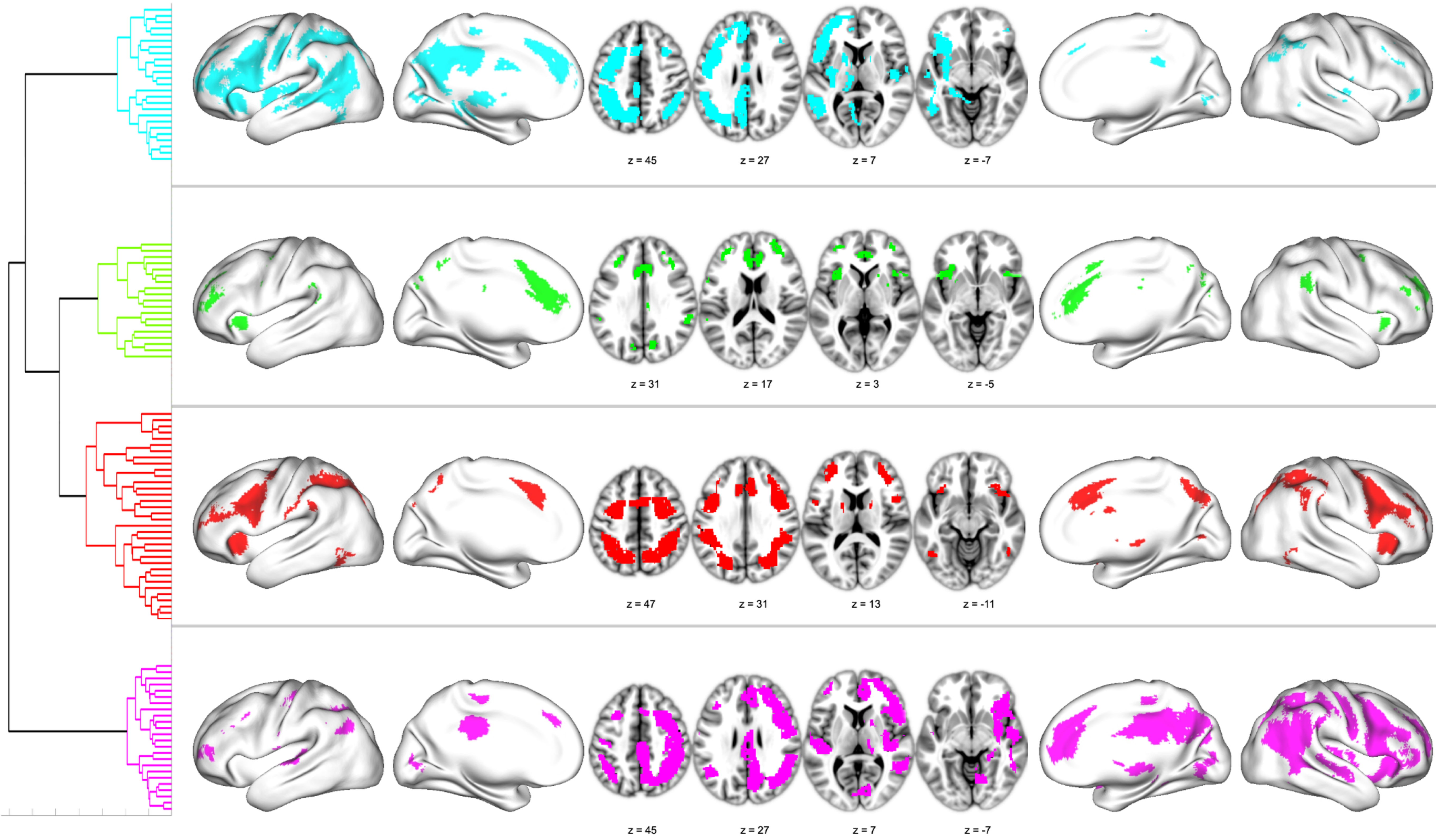

Supplemental Figure 2

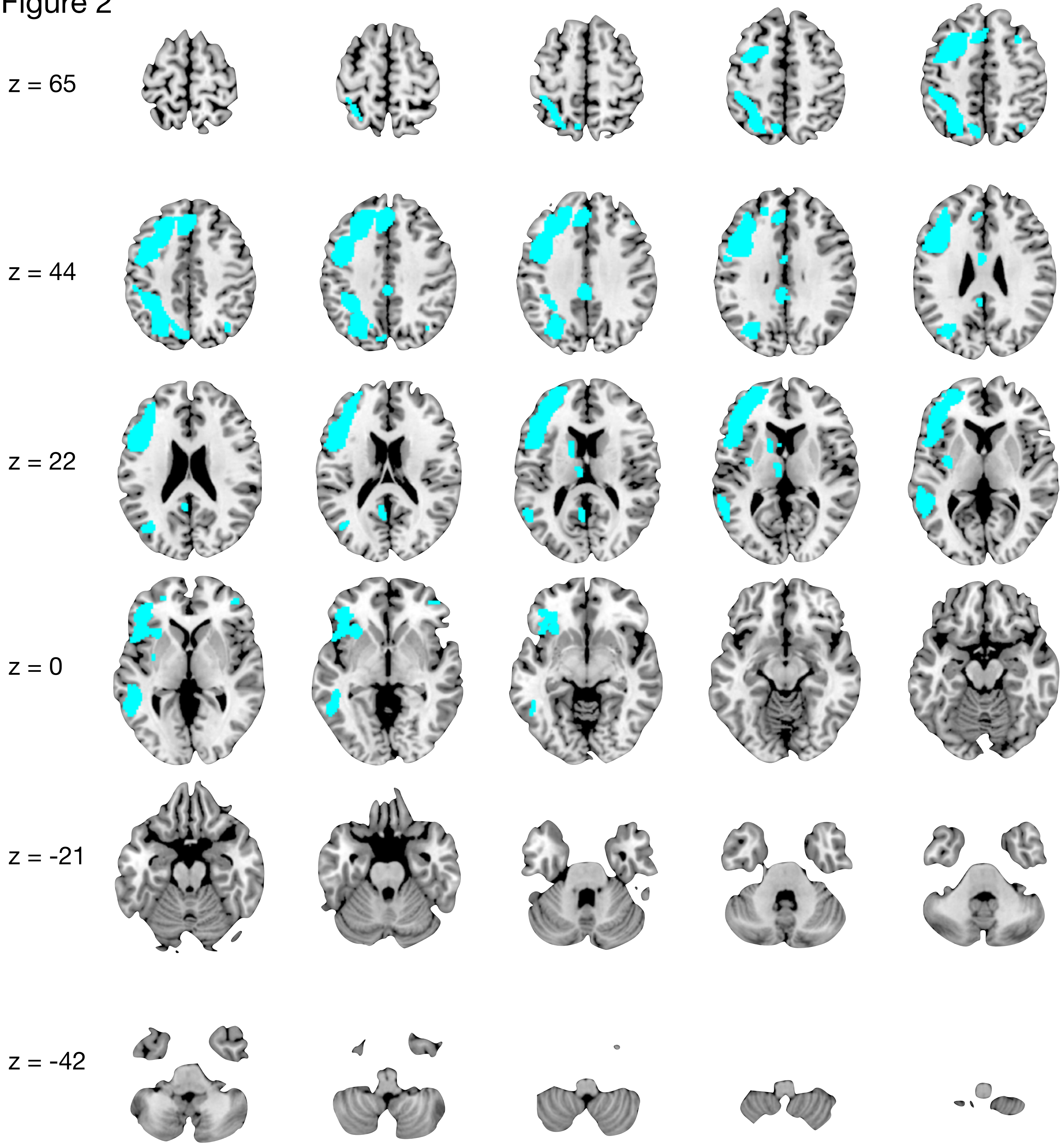

Supplemental Table 2: Cyan Cluster peak regions ($p < 0.05$)

| Cluster | Volume (mm^3) | Hemisphere | Label | x | y | z | 1-p |
|---|---|---|---|---|---|---|---|
| 1 | 35848 | L | Middle Frontal Gyrus, BA10 | -44 | 44 | 10 | 10.3 |
| | | L | Middle Frontal Gyrus, BA9 | -50 | 16 | 30 | 10 |
| | | L | Middle Frontal Gyrus, BA9 | -42 | 32 | 26 | 9.6 |
| | | L | Middle Frontal Gyrus, BA10 | -30 | 60 | 10 | 9 |
| | | L | Medial Frontal Gyrus, BA8 | -2 | 26 | 44 | 8.3 |
| | | L | Middle Frontal Gyrus, BA6 | -24 | 6 | 50 | 7.4 |
| 2 | 11144 | L | Precuneus, BA7 | -6 | -72 | 50 | 9.7 |
| | | L | Precuneus, BA7 | -24 | -70 | 50 | 9.5 |
| | | L | Precuneus, BA19 | -32 | -74 | 32 | 7.9 |
| | | L | Superior Parietal Lobule, BA7 | -30 | -52 | 60 | 5.8 |
| | | L | Inferior Parietal Lobule, BA40 | -46 | -34 | 52 | 5.3 |
| | | L | Inferior Parietal Lobule, BA40 | -32 | -42 | 46 | 5.3 |
| 3 | 2736 | L | Inferior Temporal Gyrus, BA37 | -54 | -54 | -4 | 7.4 |
| | | L | Middle Temporal Gyrus, BA22 | -60 | -38 | 4 | 5.1 |
| 4 | 1208 | L | Posterior Cingulate, BA30 | -6 | -52 | 18 | 7.5 |
| | | L | Cingulate Gyrus, BA31 | -2 | -36 | 34 | 7 |
| 5 | 312 | L | Caudate, Caudate Body | -14 | 6 | 10 | 5.5 |
| 6 | 248 | L | Claustrum | -34 | -6 | 6 | 4.9 |
| 7 | 216 | L | Thalamus, Medial Dorsal Nucleus | -6 | -14 | 10 | 5.1 |
| 8 | 184 | R | Superior Parietal Lobule, BA7 | 34 | -68 | 48 | 5.2 |
| 9 | 128 | L | Cingulate Gyrus, BA24 | -4 | -4 | 28 | 6.1 |
| 10 | 80 | R | Middle Frontal Gyrus, BA10 | 42 | 46 | -2 | 4.8 |

Supplemental Figure 3

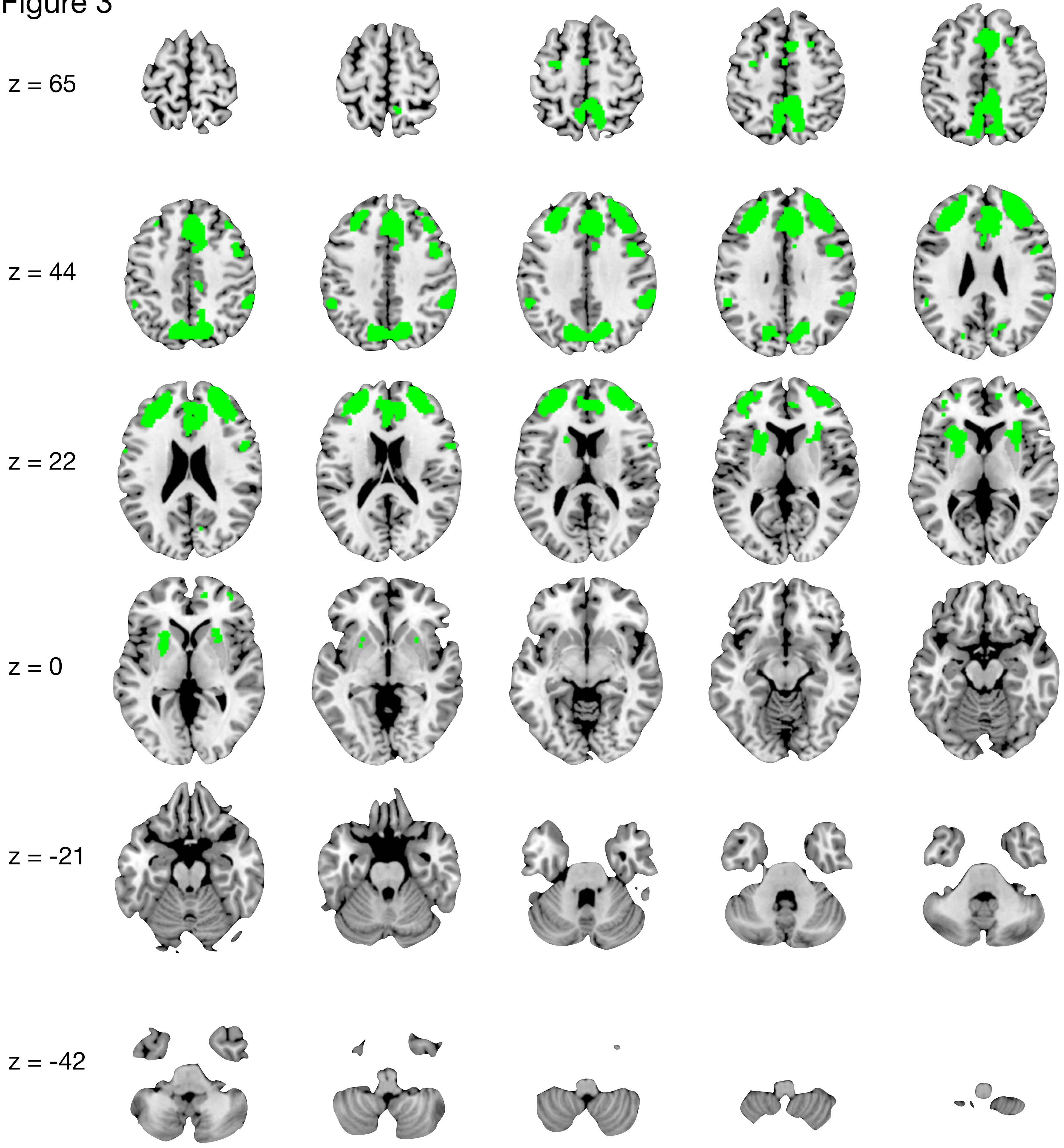

Supplemental Table 3: Green Cluster peak regions ($p < 0.05$)

| Cluster | Volume (mm^3) | Hemisphere | Label | x | y | z | 1-p |
|---|---|---|---|---|---|---|---|
| 1 | 12760 | R | Medial Frontal Gyrus, BA8 | 6 | 22 | 48 | 5.8 |
| | | R | Medial Frontal Gyrus, BA9 | 8 | 36 | 32 | 5.7 |
| | | R | Medial Frontal Gyrus, BA10 | 10 | 46 | 14 | 5.5 |
| | | L | Medial Frontal Gyrus, BA9 | -8 | 46 | 18 | 5.1 |
| | | R | Medial Frontal Gyrus, BA6 | 4 | 2 | 50 | 4.7 |
| | | L | Cingulate Gyrus, BA24 | 0 | 14 | 26 | 4.7 |
| 2 | 12136 | R | Precuneus, BA7 | 4 | -74 | 42 | 6.7 |
| | | L | Precuneus, BA7 | -14 | -76 | 42 | 6.4 |
| | | R | Paracentral Lobule, BA5 | 4 | -44 | 54 | 6.1 |
| | | R | Superior Parietal Lobule, BA7 | 12 | -62 | 56 | 5.4 |
| | | L | Precuneus, BA7 | -6 | -56 | 48 | 5.1 |
| | | R | Paracentral Lobule, BA31 | 8 | -28 | 44 | 4.8 |
| 3 | 10088 | R | Middle Frontal Gyrus, BA10 | 28 | 52 | 18 | 6.4 |
| | | R | Middle Frontal Gyrus, BA8 | 40 | 28 | 38 | 6.1 |
| | | R | Superior Frontal Gyrus, BA9 | 26 | 48 | 34 | 6 |
| 4 | 9288 | L | Superior Frontal Gyrus, BA10 | -24 | 54 | 22 | 6.6 |
| | | L | Middle Frontal Gyrus, BA9 | -38 | 28 | 30 | 6.5 |
| | | L | Middle Frontal Gyrus, BA10 | -40 | 44 | 18 | 6.4 |
| | | L | Middle Frontal Gyrus, BA9 | -28 | 40 | 36 | 5.6 |
| | | L | Superior Frontal Gyrus, BA10 | -24 | 54 | 4 | 4.8 |
| 5 | 1952 | L | Lentiform Nucleus, Putamen | -24 | 4 | 4 | 5.4 |
| | | L | Insula, BA13 | -32 | 20 | 6 | 4.8 |
| 6 | 1648 | R | Precentral Gyrus, BA6 | 44 | 2 | 36 | 5.7 |
| | | R | Inferior Frontal Gyrus, BA44 | 58 | 8 | 16 | 5.1 |
| 7 | 1560 | R | Inferior Parietal Lobule, BA40 | 52 | -44 | 40 | 6 |
| 8 | 776 | R | Claustrum | 26 | 18 | 6 | 5.1 |
| 9 | 512 | L | Inferior Parietal Lobule, BA40 | -54 | -44 | 36 | 5.5 |
| 10 | 136 | L | Precentral Gyrus, BA6 | -32 | -6 | 56 | 5 |
| 11 | 112 | R | Cingulate Gyrus, BA24 | 8 | 8 | 36 | 4.8 |
| 12 | 96 | L | Medial Frontal Gyrus, BA6 | 0 | -4 | 56 | 4.7 |
| 13 | 88 | R | Superior Frontal Gyrus, BA6 | 22 | 14 | 50 | 4.9 |

Supplemental Figure 4

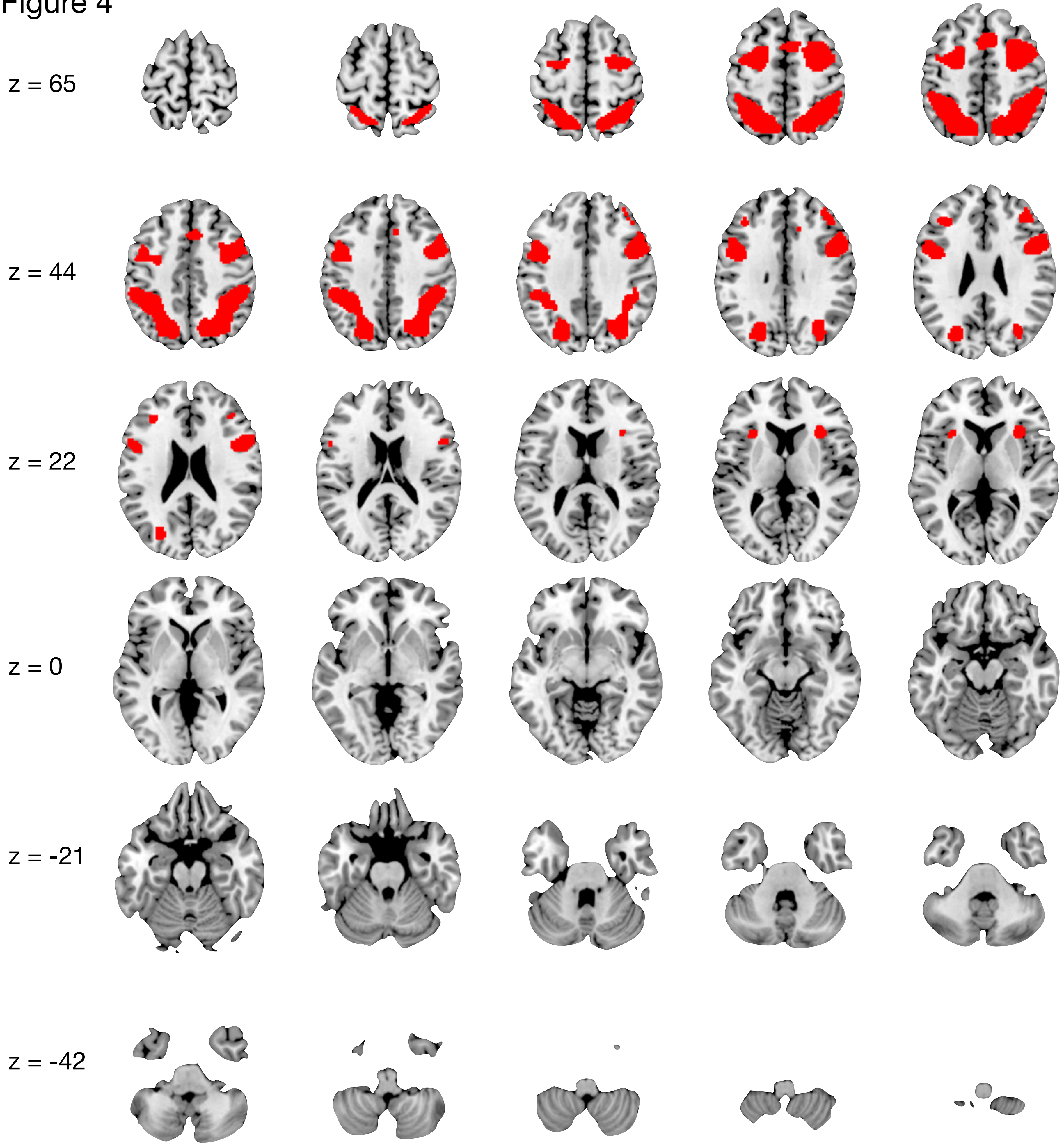

Supplemental Table 4: Red Cluster peak regions ($p < 0.05$)

| Cluster | Volume (mm^3) | Hemisphere | Label | x | y | z | 1-p |
|---|---|---|---|---|---|---|---|
| 1 | 17064 | R | Superior Parietal Lobule, BA7 | 28 | -58 | 50 | 7.3 |
| | | R | Precuneus, BA7 | 14 | -70 | 48 | 6.9 |
| | | R | Inferior Parietal Lobule, BA40 | 40 | -48 | 50 | 6.2 |
| | | R | Inferior Parietal Lobule, BA40 | 48 | -36 | 44 | 5.6 |
| 2 | 16064 | L | Superior Parietal Lobule, BA7 | -30 | -62 | 54 | 7.1 |
| | | L | Inferior Parietal Lobule, BA40 | -40 | -46 | 50 | 6.5 |
| | | L | Inferior Parietal Lobule, BA40 | -52 | -36 | 46 | 5.7 |
| 3 | 11088 | R | Sub-Gyral, BA6 | 28 | -2 | 56 | 7.9 |
| | | R | Middle Frontal Gyrus, BA8 | 52 | 8 | 38 | 6.9 |
| 4 | 6488 | L | Precentral Gyrus, BA6 | -50 | 2 | 36 | 7.1 |
| | | L | Sub-Gyral, BA6 | -28 | -4 | 54 | 6.2 |
| 5 | 1152 | L | Superior Frontal Gyrus, BA6 | 0 | 10 | 52 | 5.2 |
| 6 | 480 | R | Insula, BA13 | 32 | 18 | 10 | 4.9 |
| 7 | 472 | R | Superior Frontal Gyrus, BA9 | 38 | 38 | 30 | 4.7 |
| 8 | 304 | L | Middle Frontal Gyrus, BA9 | -34 | 32 | 24 | 4.8 |
| 9 | 104 | L | Insula, BA13 | -30 | 18 | 6 | 4.6 |

Supplemental Figure 5

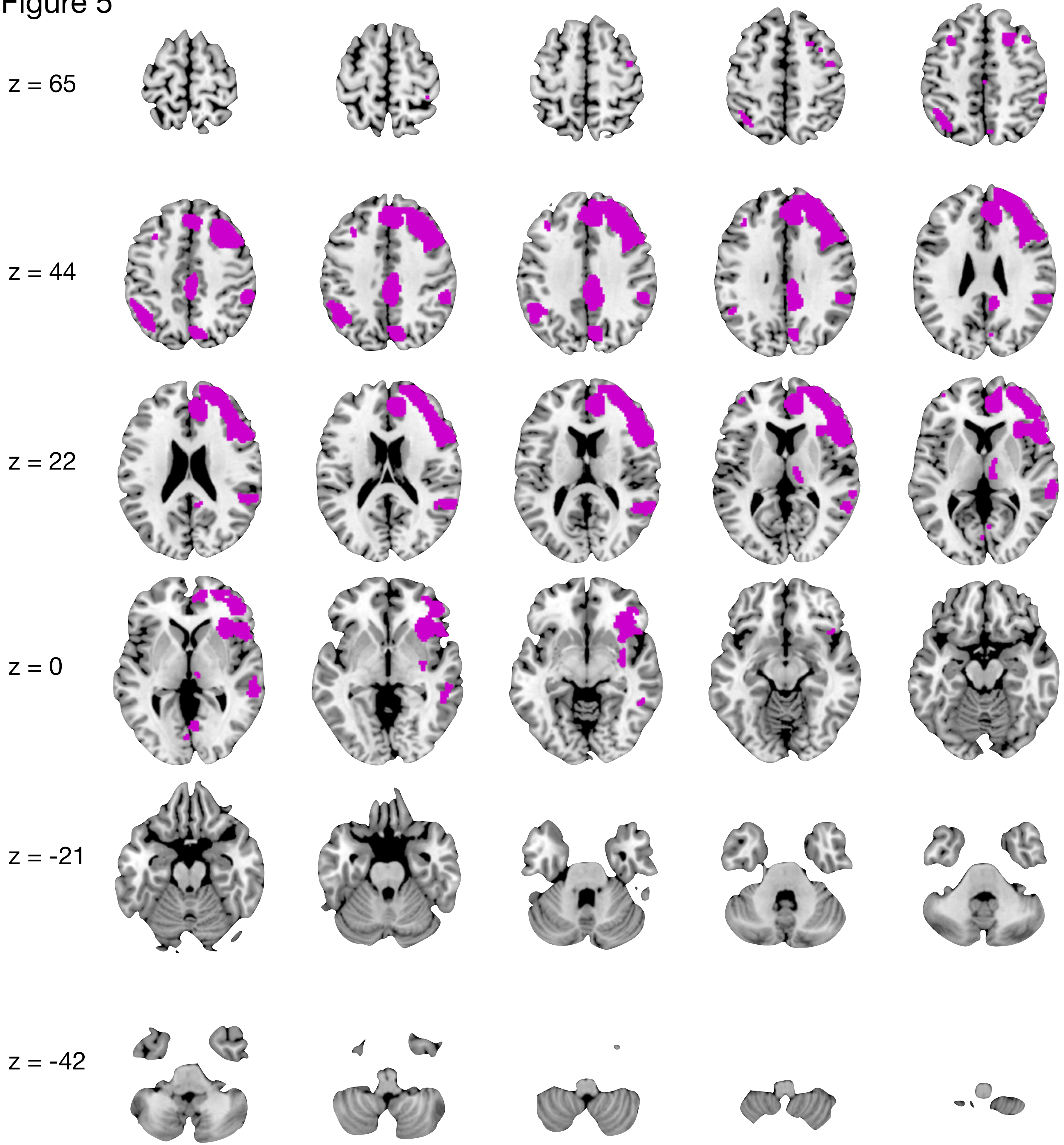

Supplemental Table 5: Purple Cluster peak regions ($p < 0.05$)

| Cluster | Volume (mm^3) | Hemisphere | Label | x | y | z | 1-p |
|---|---|---|---|---|---|---|---|
| 1 | 47528 | R | Superior Frontal Gyrus, BA10 | 24 | 62 | 16 | 10.7 |
| | | R | Middle Frontal Gyrus, BA10 | 44 | 44 | 10 | 10.5 |
| | | R | Middle Frontal Gyrus, BA9 | 36 | 44 | 28 | 9 |
| | | R | Inferior Frontal Gyrus, BA47 | 54 | 20 | 2 | 8.5 |
| | | R | Anterior Cingulate, BA32 | 10 | 38 | 20 | 8.1 |
| | | R | Insula, BA13 | 34 | 22 | 2 | 7.1 |
| 2 | 4096 | R | Cingulate Gyrus, BA31 | 4 | -42 | 34 | 9.3 |
| 3 | 3352 | R | Superior Temporal Gyrus, BA39 | 44 | -50 | 14 | 5.3 |
| | | R | Superior Temporal Gyrus, BA13 | 52 | -40 | 24 | 5.3 |
| | | R | Inferior Parietal Lobule, BA40 | 54 | -38 | 42 | 5.1 |
| 4 | 3088 | L | Inferior Parietal Lobule, BA40 | -52 | -52 | 38 | 5.6 |
| | | L | Superior Parietal Lobule, BA7 | -40 | -62 | 48 | 5.5 |
| 5 | 1656 | R | Precuneus, BA7 | 6 | -74 | 40 | 7.3 |
| 6 | 960 | R | Middle Temporal Gyrus, BA22 | 66 | -32 | 4 | 5.3 |
| | | R | Sub-Gyral, BA37 | 50 | -46 | -6 | 5.2 |
| 7 | 336 | L | Middle Frontal Gyrus, BA9 | -38 | 28 | 34 | 7 |
| | | L | Superior Frontal Gyrus, BA6 | -32 | 14 | 50 | 5.6 |
| 8 | 304 | R | Thalamus, Medial Dorsal Nucleus | 8 | -16 | 8 | 5.4 |
| 9 | 280 | R | Claustrum | 34 | -6 | -8 | 5.4 |
| 10 | 120 | R | Lingual Gyrus, BA18 | 2 | -70 | 2 | 5.1 |
| 11 | 104 | R | Precentral Gyrus, BA6 | 40 | -8 | 56 | 5.3 |

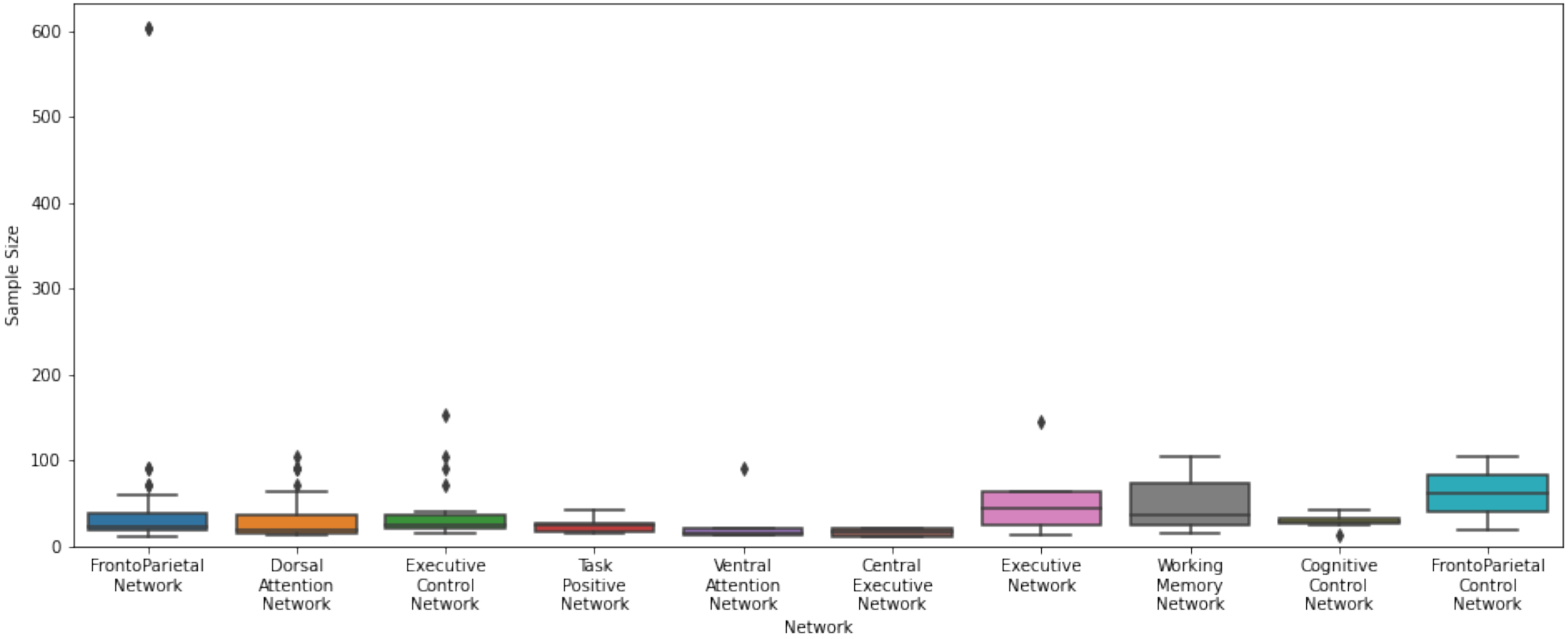

Sample Size
600
500
400
300
200
100
0
FrontoParietal Network
Dorsal Attention Network
Executive Control Network
Task Positive Network
Ventral Attention Network
Central Executive Network
Executive Network
Working Memory Network
Cognitive Control Network
FrontoParietal Control Network
Network
